# Hybridisation and herbivory fuel Amazonian tree radiations

**DOI:** 10.64898/2026.06.17.732868

**Authors:** Rowan J. Schley, Alex D. Twyford, María-José Endara, Dale L. Forrister, Akira A. Wong Sato, Carlos Reynel, James Nicholls, Graham N. Stone, Mark Blaxter, Meng Lu, Flávia Fonseca Pezzini, Caroline Howard, Thomas C. Mathers, Shane McCarthy, Johnathan Wood, Chenxi Zhou, Haroldo C de Lima, Danilo M. Neves, Maristerra R. Lemes, Luciano Paganucci de Queiroz, Phyllis D. Coley, Catherine Kidner, Kyle G. Dexter, R. Toby Pennington

## Abstract

Tropical rainforests, and Amazonia in particular, contain more tree species than anywhere else, most of which arose through rapid evolutionary radiations ^1–3^. Rapid radiations are often catalysed by ecological opportunity ^4–6^, which in rainforest trees is presented by intense insect herbivore pressure, spurring the evolution of novel plant defence chemistry to escape it ^7^. However, we do not understand how long-lived trees can adapt quickly enough to keep pace with rapidly-evolving insect herbivores. Here we show that hybridisation in rainforest trees, which was considered rare, allows exchange of gene clusters used in chemical defence against herbivore attack, facilitating rapid adaptation and diversification. Using genome sequencing for 461 individuals from the genus *Inga*, a characteristic Amazonian tree radiation, we find that regional tree communities form syngameons - networks of closely related, co-occurring species connected by gene flow. Integrating these genomes with herbivore abundance data from the same communities across the tropical Americas, we show that herbivore compositional turnover coincides with local, recurrent interspecific transfer of defence gene clusters that are retained by balancing selection, consistent with fluctuating selective pressure imposed by shifting herbivore communities. Together, our results demonstrate that hybridisation allows long-lived tropical trees to rapidly evolve chemical defences, fuelling adaptation to the relentless insect herbivory that structures the world’s most species-rich forests.

## Main

Rapid evolutionary radiations that generate species-rich groups have played a key role in assembling the most diverse biotas ^8, 9^. Neotropical rainforests are the most species-rich forests on Earth ^10^ and the Amazon rainforest is particularly emblematic, with more tree species found in a single hectare than in all of Europe ^11–13^. More than 50% of Amazonian trees belong to species-rich, evolutionarily young genera that result from rapid evolutionary radiation ^1–3^. Such rapid radiations require ecological opportunity for diverging populations to occupy new fitness optima ^4–6^, which in tropical rainforests can be provided by biotic interactions ^14^. In particular, relentless pressure from insect herbivores ^15^ imposes potent density-dependent mortality on seedlings and saplings ^16–18^. The high herbivory pressure of tropical forests has led to rapid diversification of anti-herbivore defences alongside speciation in rainforest trees ^7^, facilitating co-existence among many closely-related tree species in local rainforest communities ^19, 20^, because differing defences mean fewer shared herbivores. Yet, it remains unclear how rainforest trees with comparatively long generation times ^21^ can generate sufficient genetic variation through mutation alone to keep pace with an evolutionary arms race against rapidly reproducing insect herbivores^22^.

Hybridisation is one mechanism that can facilitate rapid adaptation, particularly in syngameons – groups of closely-related, co-occurring species that recurrently interbreed at low frequency, forming networks of hybridising species ^23–25^. In syngameons, which have been documented in groups such as cichlid fishes and temperate oaks, it has been hypothesised that occasional hybridisation promotes ecological adaptation and the emergence of novel phenotypes by redistributing and recombining genetic variation via introgression (the transfer of genetic material between species through hybridisation and backcrossing) ^25, 26^. Introgression of defence chemistry loci within syngameons provides a plausible mechanism by which tropical trees could rapidly adapt to insect herbivores. However, this hypothesis remains untested, as hybridisation was historically considered extremely rare in tropical trees ^27^ and was consequently overlooked as an evolutionary force (reviewed in ^28, 29^).

Here, we generate and analyse a unique dataset comprising extensive whole-genome sequencing, plant chemistry and insect herbivore data to test the syngameon model of hybridisation in tropical rainforest trees. We investigate whether hybridisation in syngameons generates functional genetic diversity, fuelling rapid evolutionary radiations, focusing on the neotropical tree genus *Inga* (Fabaceae). *Inga* typifies the species-rich tree radiations that generated most Amazonian tree diversity ^1, 30^ because of its ubiquity in neotropical rainforests ^31^, its species-richness (*ca.* 300 species diversified in the last 7*-*10 Ma ^32^), its high levels of co-existence in local communities (up to 19 *Inga* species in 1 ha ^33^), and its diverse anti-herbivore defence chemistry ^34–36^. There is also emerging evidence of hybridisation in *Inga* ^37, 38^, likely the result of overlapping phenology and pollination syndromes ^39^ (Figure 1a), rendering it an ideal model clade *(c.f.* ^40^) with which to understand the role of hybridisation in generating tropical tree diversity. This framework makes syngameons particularly relevant for understanding how ecological interactions, such as herbivory, can shape adaptive radiations in species-rich tropical forests. Our results reveal that hybridisation in local syngameons likely fuels adaptation by shuffling chemical defence genes among species, generating functional variation in herbivore defence. This rapid, ‘combinatorial’ ^41^ evolution of novel defences allows trees to keep pace in the evolutionary arms race with their herbivores, catalysing the assembly of the most species-rich tree floras on Earth.

**Figure 1:**
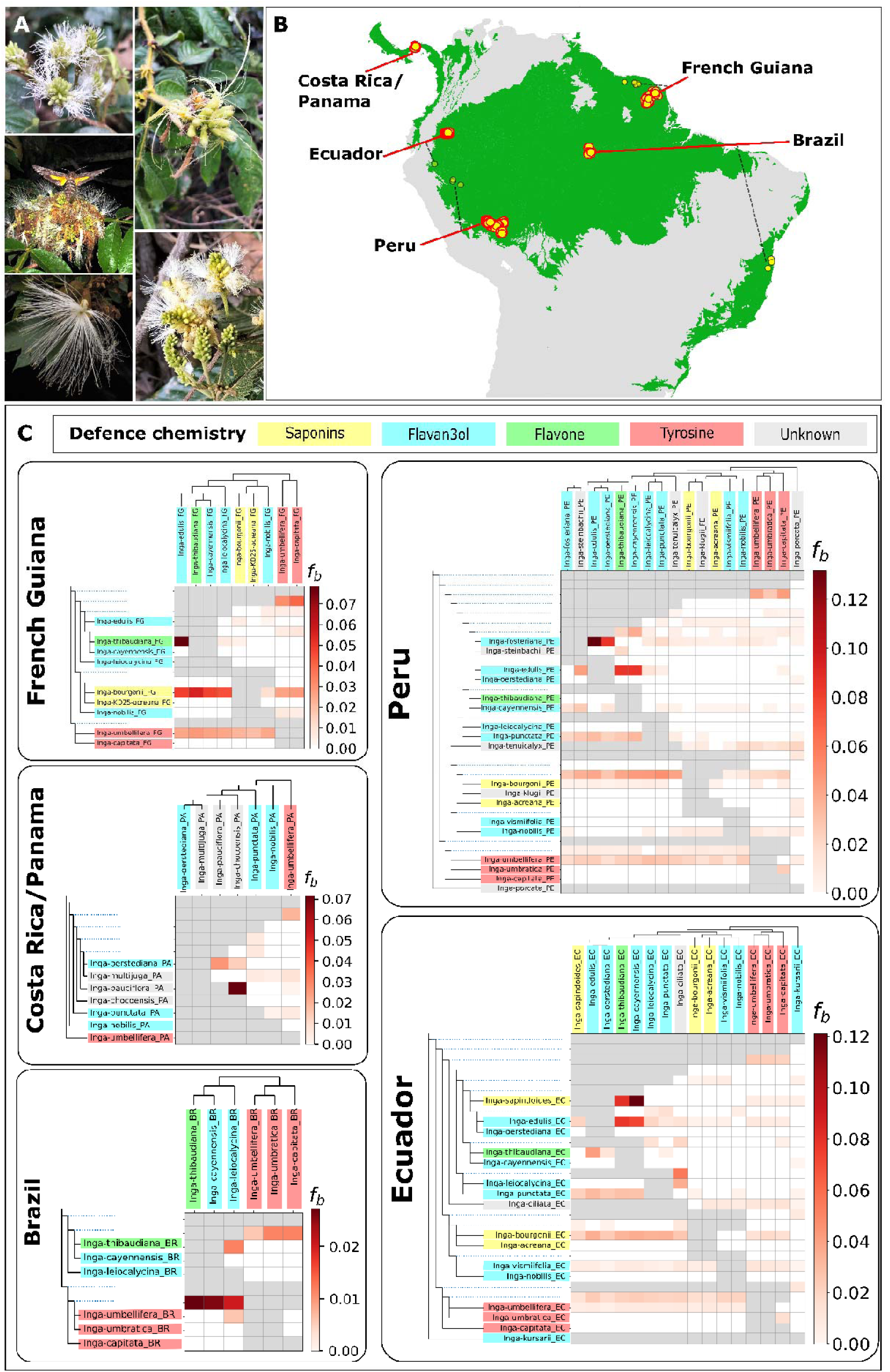
Introgression occurs in regional *Inga* communities across the tropical Americas. **a**) *Inga* inflorescences, demonstrating their similarity and generalist pollination syndrome. Clockwise from top left: *Inga auristellae* (© Rowan Schley), *I. setosa* (© Rowan Schley), *I. lineata* (© Toby Pennington), *I. sessilis* (© Toby Pennington) and *I. edulis* being visited by a hawkmoth (*Erinnyiss* sp.), one of several common *Inga* pollinators (© Toby Pennington). **(B)** Map of South and Central America showing sampled regional *Inga* communities for each tropical American country. Points represent sampling localities, and are plotted with jitter where localities were identical. Large points with a bold red outline and red arrows represent collections from core research stations in five regions: Barro Colorado Island (Panama), Nouragues (French Guiana), Reserva Ducke and PDBFF in Manaus (Brazil), Yasuní (Ecuador) and Madre de Dios (Peru). Small points with black outlines and black hashed arrows represent other collections made to ensure accurate representation of genetic diversity. Green shading marks the distribution of tropical rainforest in Central and South America, modified from ^29^. **(C)** Heatmaps of ‘F_branch_’ scores, a summary of introgression between species (x axis) and branches (y axis), here plotted for each regional *Inga* community. The colour of each square in each heat map signifies the amount of excess allele sharing due to introgression (f_b_; dark red = high estimate). In each plot, the phylogenetic tree of the species found in a community is displayed in an ‘expanded’ form along the y axis, so that all branches (including internal ones, marked as dotted lines), correspond to a row in the heatmap. The same tree is represented in an unexpanded format on the x axis, such that each tip corresponds to a column in the heatmap. Grey boxes in the heatmap indicate F_branch_ tests that were not possible to perform due to the topology of the tree, e.g. between sister species. The main class of chemical defence (where known) used by each species is displayed as a coloured box around the taxon name, after ^7^, defined in the legend at the top of Figure 1c.

## Rainforest trees form local syngameons

Here, we test for recurrent, geographically-structured hybridisation between multiple species, as expected in syngameons. Our whole-genome resequencing of 461 individuals, comprising 24 *Inga* species from five regional rainforest communities spanning the tropical Americas, unveils introgression between co-occurring species consistent with the existence of syngameons.

Genome-wide tests for introgression (F_branch_ ^42^) in regional *Inga* communities recovered up to 12% introgressed ancestry (Figure 1; Fig. S1a-b, Fig. S2a-e and S12, S3a-f; Extended Data). We found excess shared ancestry in multiple species pairs within regional communities, with several involved in hybridising species networks (i.e. syngameons) comprising more than one other species (e.g. between *Inga umbellifera* and three other species in Peru; Figure 1; Fig. S1a-b, Fig. S4, Extended Data). We also inferred widespread population-level phylogenetic discordance; for the plastid genome, accessions frequently grouped by geography rather than species, in contrast to the nuclear genome that recovered morphologically-diagnosable *Inga* species (Fig. S5 and S13, Extended Data). Such a pattern of phylogenetic conflict suggests that hybridisation is infrequent enough for species to maintain their integrity, and that it occurs locally within *Inga* communities leading to regional plastid capture, both of which are hallmarks of syngameons in plants ^43–45^.

If co-occurring species form syngameons, we expect a comparatively small portion of the genome (likely underlying reproductive isolation) to maintain species divergence over long periods of time. We used block-wise demographic inference ^46^ to estimate the proportion of genomic windows acting as barriers to gene flow over time, using a subset of eight closely related *Inga* species pairs from the same regional communities (selected to ensure a mix of chemical defence strategies and phylogenetic distances between species pairs, further detailed in Table S3 and Methods, Extended Data). Our analyses indicate that long-term barriers to gene flow comprise between 0.08% - 5.45% of genomic windows in these eight *Inga* species pairs (Table S5), suggesting that only a limited portion of the genome has contributed to restricting gene flow over time, even among the most evolutionarily divergent species pairs we studied (e.g. *Inga capitata* and *I. tenuicalyx*). Our estimates of the proportion of the genome acting as a barrier to gene flow in *Inga* are similar to those from other syngameon-forming genera (e.g. 0.78% of the genome in *Heliconius* butterflies ^46^; 1.99% of the genome in *Populus* trees ^47^), and are substantially lower than estimates for taxa not known to form syngameons (e.g. 23.08% of the genome in *Brenthis* butterflies ^48^).

## Defence genes recurrently introgress

Having established that syngameons occur in regional tropical rainforest tree communities, we tested whether recurrent introgression among non-sister species transfers functional genetic variation that promotes ecological diversification, as predicted by the syngameon model of adaptive radiation ^45, 49^. We focus on plant defence chemistry, where new phenotypes arise largely through reassortment of existing chemical subunits into new combinations (“Lego chemistry” ^7, 50^) rather than *de-novo* innovation, which should be more rare. The model is most compelling in syngameons, because introgression provides access to defence-related gene modules from multiple species that can be recombined into new phenotypes.

Biosynthetic gene clusters (BGCs), usually encompassing several dozen co-located genes associated with a certain biosynthetic pathway, play a disproportionately important role in secondary metabolism ^51^. Thus, we used PlantiSMASH ^52^ to identify 32 BGCs underlying key pathways for defence chemical diversity in *Inga* ^35^ (Table S6). Next, we identified BGCs that experience elevated introgression based on *f_dM_*scores ^53^, relative to the genome-wide average. We recovered 19 BGCs that experienced recurrent introgression between different, co-occurring non-sister *Inga* species, suggesting independent events of allele sharing between multiple species pairs (Table S7; species selection criteria for *f_dM_* described in Table S3, Extended Data). In some cases, introgression occurred in the same gene, and in others introgression occurred in different gene blocks within the same BGC (Figure 2; Table S7, Extended Data) – however, in all cases introgression involved different SNPs, suggesting multiple independent introgression events. The most notable examples were for BGCs involved in the production of flavonoids and terpenes, key anti-herbivore defences in *Inga* ^7^. This included the “*DIOX_N Glycosyltransferase 1”* and “*2OG-FeII_Oxy, DIOX_N, p450”* BGCs (both associated with modification of flavonoids ^54–56^), the “*Bet v1 Transferase”* BGC (involved in regulation of flavonoids and other defence chemicals ^57, 58^) and the “*Terpene Synthase”* BGCs (involved in the production of terpenes ^59^). Moreover, the highest *f_dM_* score we recovered across any *Inga* genome was within a BGC containing *DAHP synthase* and 10 associated genes (*f_dM_*= 0.71, where a score of 1 indicates total allele sharing; Figure 2; Table S7, Extended Data). *DAHP synthase* catalyses the start of the shikimate pathway, a crucial step in the biosynthesis of most defence chemicals in *Inga* ^35^.

**Figure 2:**
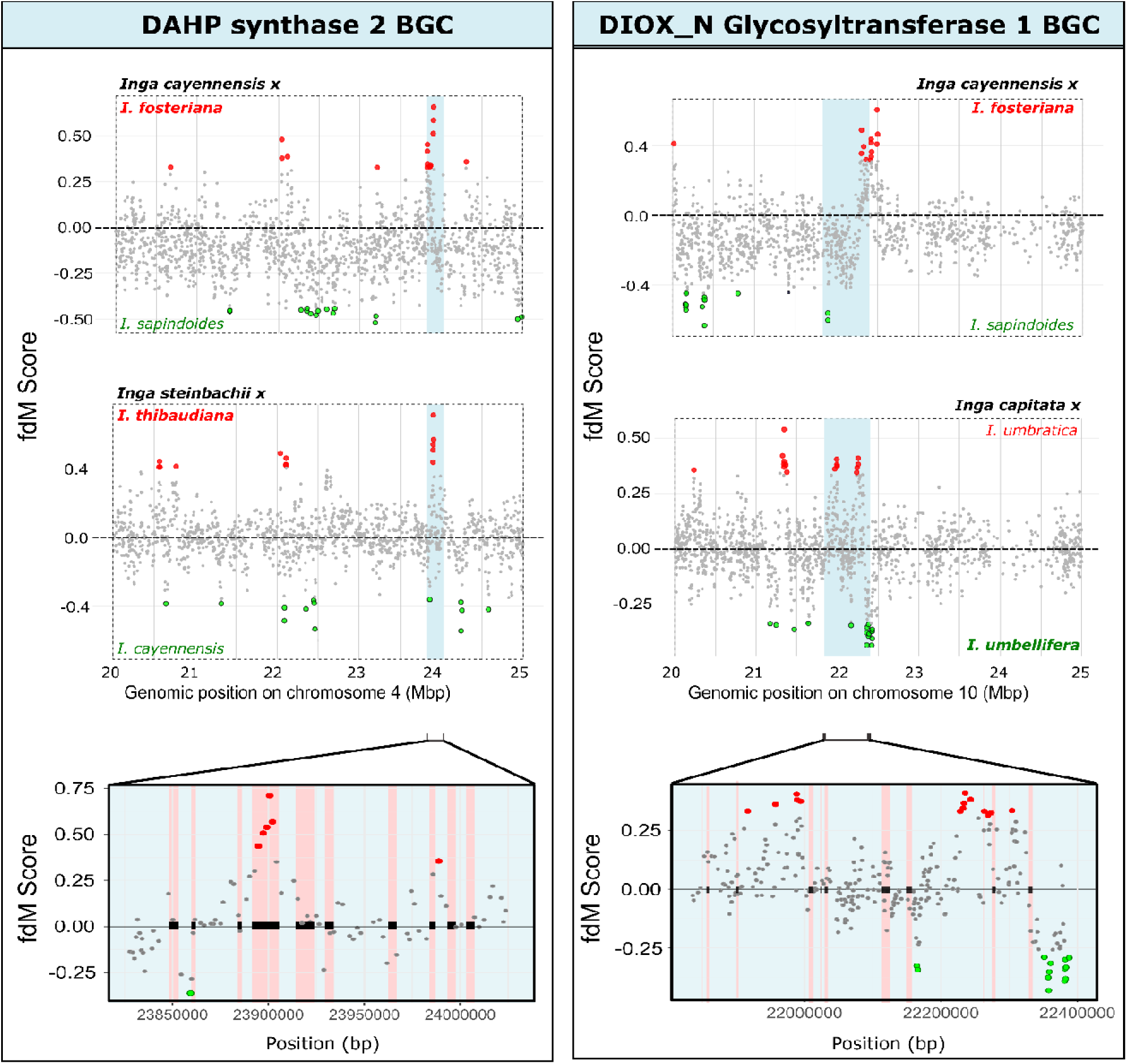
Recurrent introgression of chemical defence genes between different co-occurring *Inga* species. Recurrent introgression events occurring in the same BGCs within different *Inga* species that co-occur in local syngameons. Introgression scores (*f_dM_*) in genomic windows surrounding the “*DAHP synthase (p450)*” and “*DIOX_N Glycosyltransferase*” BGCs on chromosomes four and ten, respectively. The extent of the BGC is marked with a blue shaded box. Taxa used for each test are indicated in each plot, with P3 taxa in black above the plot, P1 taxa in green and P2 taxa in red. Positive *f_dM_* values (red outliers) indicate introgression between taxon P3 and P2, while negative scores (green outliers) indicate introgression between taxon P3 and P1. The species pair with the highest introgression scores (i.e. P3 x P2 or P3 x P1) are emboldened in each *f_dM_* plot. Outliers are first and 99^th^ percentile values. The bottom two panels indicate a zoomed in view of introgression scores within the BGC of the corresponding column: the left panel corresponds to introgression scores for the *Inga steinbachii/I.thibaudiana/I. cayennensis* taxon trio, while the panel on the left corresponds to the introgression scores for the *Inga capitata*/*I.umbratica/I.umbellifera* taxon trio. In these bottom two panels, the position of gene regions within each BGC are marked by black boxes with red background shading, and points are coloured in the same way as in the top panel, indicating which species pair introgression scores were recovered for.

Our results suggest that new combinations of chemical precursors may arise through introgression of different gene modules, particularly for flavonoids. It is well established that simple flavonoid molecules are embellished through polymerisation and addition of R-groups, meaning dozens of unique polymers can arise from one simple precursor ^7^, suggesting that re-assembly of biosynthetic genes by introgression could rapidly result in novel defence chemistry. Overall, novel genetic diversity introduced into such crucial biosynthetic pathways is expected to have a significant effect on downstream compounds ^60^, consistent with the ‘Lego chemistry’ model. As such, our results suggest that introgression in local syngameons can furnish rainforest tree species with novel genetic diversity in key regions of the genome that underlie chemical defence.

## Defence genes introgress as modules

Having established that defence-related loci introgress recurrently among co-occurring species, we next ask whether these loci persist as linked genomic ‘modules’ following introgression. To address this, we used our per-window estimates of introgression across the genome to identify 15 candidate BGCs (of the 32 BGCs identified by PlantiSMASH) with exceptionally high estimates of introgression in 23 *Inga* species pairs, totalling 88 BGC/species pair combinations. We then estimated the longest contiguous block of introgressed variation (i.e. regions containing the same, incongruent genealogy) in each candidate BGC for each introgressing species pair (Table S8; Fig. S6, Extended Data).

Introgressed blocks spanned between 0.33% - 69.25% of their respective candidate BGCs across all 88 analyses, ranging between 1 - 249 kilobases (Kb) in length. The largest block of introgressed ancestry was recovered between *Inga chocoensis* and *I. pauciflora* in Central America, for the ‘*Terpene Synthase*’ BGC (containing eight genes; Table S8, Fig. S6, Extended Data). This introgressed region was significantly larger than a random draw of introgressed regions on the same chromosome (Table S8, Extended Data). We found that BGCs relating to defence chemical regulation (e.g. the *“Beta_v1_transferase”* BGC) were prone to introgression as longer clusters than other BGC classes (between 145-220 Kb; containing two genes; up to 55% of original BGC length), most often between recently divergent *Inga* species ^37^ such as *I. steinbachii* and *I. cayenennsis* in Peru (Table S8, Fig. S6, Extended Data). Notably, while some BGCs underlying flavonoid modification (e.g. the *“DIOX_N Glycosyltransferase 1”* and “*2OG-FeII_Oxy, DIOX_N, p450”* BGCs) were retained as large blocks (up to 189Kb) following introgression between multiple *Inga* species pairs, most introgressed variation in flavonoid BGCs persisted only as smaller blocks (2.14% - 23% of the original BGC length, mostly with 1-3 genes; Table S8, Fig. S6, Extended Data).

Our results suggest that most BGCs are retained as modular ‘building blocks’ of a few hundred Kb following introgression, with most introgressed regions containing around two genes (Table S8, Fig. S6, Extended Data). Thus, our results provide further evidence for the ‘Lego chemistry’ model of defence evolution in *Inga* ^7, 50^, suggesting that introgression and within-BGC recombination play an important role in reassortment of genomic ‘building blocks’ that underlie defence chemistry.

The varying proportions of introgressed variation between different BGC classes may be explained by their genomic architectures, which alter how rapidly recombination breaks down introgressed blocks ^61, 62^. For example, the most contiguous introgressed BGCs we found involved clusters of up to 10 genes underlying embellishment of terpenes (*Terpene synthases*) and regulation of defence chemicals (*Bet V1 transferases*). Genes associated with biosynthesis of terpenes have been shown to be co-located in large, contiguous clusters ^63^. Similarly, *Bet V1 transferase* is known to occur in compact, tightly-linked clusters with low-recombination that underlie defence chemistry (e.g. the BIA alkaloid gene cluster in poppies (*Papaver*) ^63–65^). In contrast, flavonoid genes have been demonstrated to exhibit a more dispersed genomic architecture, often being loosely linked ^63^. The broad variation in the size of introgressed blocks that we observed for flavonoid BGCs (Fig. S6, Extended Data), suggests re-assortment of loosely-linked genes by introgression, and could explain how so much variation in flavonoid molecules can be produced from simple precursors among *Inga* species ^7^. Overall, our results echo recent work demonstrating how introgression of modular gene clusters underlies diversification of ecological traits during other rapid adaptive radiations, such as in the phenotypically diverse lake Victoria cichlid fish radiation ^66^.

## Selection maintains introgressed defence

Rare chemical-defence variants should be advantageous in tree populations because local herbivores are less likely to possess counter-adaptations - we therefore predict balancing selection, via negative frequency dependence ^16, 18, 67^, will maintain introgressed defence gene variants after they enter recipient populations. Consistent with this, our analyses recovered balancing selection in multiple introgressed chemical defence gene blocks (Table S8 – S10, Extended Data). Elevated balancing selection was evident in many of the same BGCs as were subject to introgression, namely those involved in the modification of flavonoids (the “*DIOX_N Glycosyltransferase 1”* and “*2OG-FeII_Oxy, DIOX_N, p450”* BGCs) and regulation of defence chemicals (“*Methyltransf_2,UDPGT_2”* BGC) alongside the production of terpenes (“*Terpene Synthase”* BGC) (Table S8 - S10, Extended Data). Recurrent balancing selection on introgressed BGCs was most frequent between a suite of *Inga* species, almost all of which were from Peru and Ecuador (Fig. S7, Table S8 – S10, Extended Data). The longest tract of introgressed variation in which we observed balancing selection was between *Inga klugii* and *Inga tenuicalyx* in Peru (109Kb in length, within the *2OG-FeII_Oxy, DIOX_N, p450 BGC* which is associated with flavonoid modification) (Fig. S8, Table S8–S10, Extended Data). Notably, *I. tenuicalyx* also showed evidence of introgression in recent phylogenomic work ^37^. Genomic windows with both elevated introgression and balancing selection were not restricted to genomic regions with elevated repeat density or SNP density, meaning that our results are likely not the result of methodological artefacts (Fig. S9 and S14, Extended Data).

Our results are congruent with adaptive introgression, in particular where introgressed variants are maintained by balancing selection ^68, 69^. We found recurrent introgression in the same BGCs across multiple co-occurring species, suggesting that syngameons allow assembly of shared ‘pools’ of variation among networks of different species, which is then subject to balancing selection. Our results are thus analogous to wing-colour mimicry genes in *Heliconius* butterflies, where introgression facilitates convergent adaptation to local mimicry rings through frequency-dependent selection on introgressed variation, allowing evasion of local predators ^70^. Loci under balancing selection are often among the last to stop introgressing as species diverge ^71^, often in regions of the genome subject to negative frequency-dependent selection (e.g. in *Arabidopsis* ^72^), and in particular those relating to defence against natural enemies (e.g. in *Helianthus* ^73^). Indeed, it has been suggested that regions subject to negative density-dependence and balancing selection can promote the evolution of porous species boundaries, due to the fitness advantages brought by occasional gene flow ^74^.

## Herbivory shapes selection on defence

If insect herbivory is a major factor influencing selection on rare, introgressed genetic variation underlying chemical defence, a necessary precondition is high turnover in herbivore community composition - if herbivore species composition was homogeneous, then there would be less selection for diversity in defence (e.g. ^75^). Furthermore, we expect balancing selection in genes underlying chemical defence to be associated with variation in herbivore communities ^76^.

Our dataset of DNA-barcoded *Inga* herbivores, comprising 5,740 herbivores from 522 lineages in 13 insect families (Fig. S10, Extended Data), shows that herbivore species turnover was explained by both regional community (Panama, French Guiana, Ecuador, Peru, F = 4.175; *P* = 0.00001, n = 126) and *Inga* host species (F = 1.112; *P* = 0.00002, n = 126), with many herbivores only found in a single region (Figure 3a; Fig. S11, Table S11, Extended Data).

**Figure 3.**
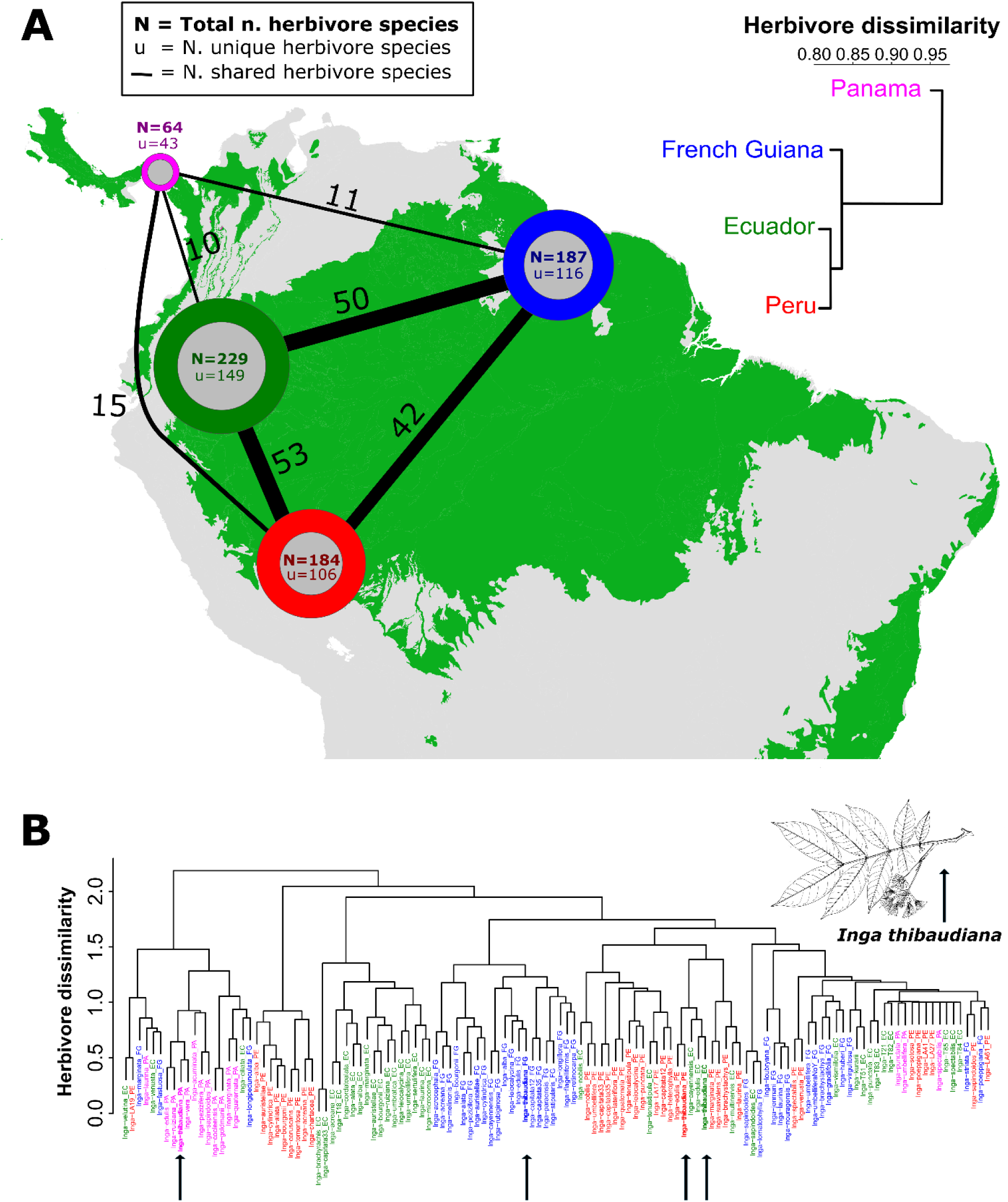
: *Inga* herbivore communities differ between neotropical rainforest regions and host species. **a**) Map of South and Central America, with circles showing the total number of *Inga* herbivore species in a regional community (N; coloured circle size) alongside the number of unique *Inga* herbivore species in a regional community (U; inset grey circle size) and the number of shared *Inga* herbivore species between each regional community (line width). Herbivore species counts in this panel represent all herbivores associated with any *Inga* species in each regional community. Herbivore communities are coloured by region – Panama (pink), French Guiana (blue), Ecuador (green) and Peru (blue). The dendrogram to the right of the map shows herbivore community dissimilarity (Bray-Curtis distances) between the four focal communities, coloured by community as above. Green shading marks the distribution of tropical rainforest in Central and South America, modified from ^29^. **(B)** Dendrogram of herbivore community dissimilarity (Bray-Curtis distances) for herbivores found on specific *Inga* species in regional communities. Tip labels represent the collection of herbivore species found on a specific *Inga* species in a regional community (e.g. *Inga_thibaudiana_EC* represents all herbivore species found on *I. thibaudiana* in Ecuador). Tip labels are coloured by regional community, as above. Black arrows mark positions of herbivore communities found on the exemplar taxon *Inga thibaudiana* (illustrated, modified from ^31^)*. Inga thibaudiana* is present in all regional communities and demonstrates the pervasive pattern of species-specific collections of herbivores clustering by regional community rather than host species.

Hierarchical clustering of herbivore assemblages grouped Amazonian regions (French Guiana, Ecuador and Peru; Figure 3a) separate from Panama. Population level analyses showed herbivore assemblages were often more similar within the same regional community than on the same *Inga* host species from different regions (Figure 3b). Our results also show strong phylogenetic turnover in herbivores between sites and *Inga* host species, most prominently in three Lepidopteran families (Gelechiidae, Noctuidae, Riodinidae) and sawflies (Argidae) (P < 0.05; Table S12, Extended Data). Together, these results demonstrate strong differences in herbivore community composition, both between regions and within regional *Inga* communities.

Furthermore, we found a significant correlation between herbivore community composition and balancing selection profiles among different local *Inga* populations, both within the plantiSMASH BGCs (R=0.213, *P=0.020,* n = 22) and within flavonoid genes specifically (R=0.182, *P=0.044,* n = 22). Balancing selection in flavonoid genes was particularly correlated with phylogenetic turnover in Riodinid butterflies (R=0.257, *P=0.030*, n = 19) (Table S13-S14, Extended Data). These results parallel the recurrent introgression and balancing selection we found in these genes (Table S7-S10, Extended Data).

Our results suggest that balancing selection in regions of the genome relating to defence chemistry correlates with local herbivore community composition, consistent with the influence of negative density-dependent selection on defences among different herbivore communities (as in the herbs *Boechera* ^77^ and *Arabidopsis* ^78^). *Inga* seeds disperse effectively, such that the entire Amazon basin acts as a metacommunity for *Inga* ^79, 80^. This means that *Inga* species frequently encounter different herbivore communities ^81, 82^, which play a principal role in structuring local tree communities ^80^, providing opportunity for rapid adaptation fuelled by introgression. We found several cases in which introgression may have influenced adaptation. For example, *Inga umbratica* and *Inga nobilis* are primarily attacked by larvae of the metalmark butterflies (Riodinidae), and both *Inga* species invest in flavonoids, which are toxic to Lepidoptera ^7, 83^. In co-occurring populations of these species from the Peruvian Amazon, we recovered elevated introgression and balancing selection in the *“DIOX_N, Glycos_transf_1, adh_short, adh_short_C2”* BGC, which is associated with flavonoid production (Tables S6 – S8, Extended Data). This BGC, along with several others in which we found introgression, are involved in final modification of flavonoid compounds. Together, this suggests that comparatively little introgression may have significant effects on final embellishment of defence compounds in *Inga*, which then provide substrate for selection and adaptation to new herbivore communities.

## Conclusions

We provide strong evidence for the ‘syngameon’ model of adaptive radiation in fuelling ecological diversification within the Amazonian tree flora. Our results at a continental scale draw parallels with iconic adaptive radiations in insular systems that were fuelled by hybridising syngameons, such as Lake Victoria’s cichlid fishes ^25^, Hawaiian silverswords ^84^, Darwin’s finches ^85^, and *Anolis* lizards ^49^. Most significantly, we provide the first evidence that introgression in local syngameons may provide locally-adapted defences for tropical plants to evade local insect herbivores. Adaptation fuelled by introgression is rendered even more likely in *Inga* because chemical defences are expected to evolve via the ‘Lego chemistry’ model, where combinatorial addition of chemical embellishments to existing compounds generates diverse new defence chemistries ^7^. We suggest that such a modular, combinatorial architecture of trait evolution may be common in adaptive radiations more generally.

In all, we show that occasional hybridisation in syngameons generates diversity in chemical defence, which is well-known to influence species divergence ^7^ and facilitate ecological coexistence ^36^ in tropical rainforest trees, enabling them to keep pace with rapidly shifting herbivore communities. The overlapping phenology and pollination strategies typical of many rainforest tree species ^39^, alongside their low conspecific density, the predominance of outcrossing mating systems ^86^ and high levels of coexistence between closely-related species ^36^, mean that hybridisation in syngameons is likely to occur widely. Thus, syngameons may have catalysed the diversification of many rainforest tree genera, and fuelled the assembly of the most species-rich tree floras on Earth ^16–18^. Moreover, our results suggest that syngameons are an important source of genetic diversity in changing biotic environments – resilience against herbivory in *Inga* is of particular societal relevance, as *Inga* is the most widely cultivated tree for agroforestry and ecosystem restoration in the tropical Americas ^31^.

## Methods

### Sampling

We sampled 461 *Inga* accessions, comprising 24 species, from many of the same study sites (hereafter ‘regional communities’) across Latin America. These regional communities included field stations and permanent plots from Central America (Panama, Costa Rica: 3 sites), western Amazonia (Ecuador, Peru: 19 sites), eastern Amazonia (French Guiana, Guyana: 8 sites), Central Amazonia (Brazil: 4 sites) and the Atlantic Forest (Brazil: 3 sites). As outgroups we also sampled one accession each from five species of *Zygia* (the sister group to *Inga*), as well as one accession each from three *Abarema* species and one *Albizia* species, which are more distantly related to *Inga*.

Our sampling was based on a list of all accepted *Inga* species compiled using the World Checklist of Vascular Plants ^87^ (as of June 2021) and a monograph of *Inga* ^31^. All accessions were collected from the silica gel-dried leaf collections held at the Royal Botanic Garden Edinburgh (RBGE). Species were chosen based on whether they showed evidence of introgression in ^37^, while ensuring that we sampled both abundant, widespread species as well as rarer, range-restricted species (based on ^31^). We also made sure to sample species representing the range of chemical defence strategies in *Inga* (based on ^7^) and that species were sampled from across the *Inga* phylogeny (based on ^37, 88, 89^). Polyploidy is uncommon in *Inga*, and so we only sampled diploid species in order to simplify downstream analysis ^90–92^. The list of sampled accessions, alongside accession information, is held in Table S1, Extended Data.

### Whole-genome resequencing, read mapping and variant filtering

For each sample we extracted DNA from 20 mg of dried leaf material with the DNeasy Plant Mini Kit (Qiagen, Hilden, Germany). DNA library preparation and sequencing were carried out by the University of Exeter sequencing service (Exeter, UK) following the NEBnext Ultra II FS protocol (New England Biolabs, Ipswich, MA, USA). Libraries were sequenced to a mean depth of 20x using the NovaSeq 6000 platform (Illumina, San Diego, CA, USA ) with paired-end 150bp runs on S4 flow cells.

Sequence data were filtered using *fastp* v0.23.4 ^93^ to remove reads with quality scores below 22 (*-q 22*) and reads less than 75 bp long (*-l 75*), while also detecting and removing adapters *(--detect_adapter_for_pe*), allowing base correction in overlapping reads (*-c*) and trimming poly-G tails from read ends (*-g*). We used *FastQC* v0.11.9 ^94^ and *MultiQC v1.19* ^95^ to assess read quality before and after filtering. Here and throughout, all analyses were conducted on the UK Crop Diversity Bioinformatics HPC Resource.

We mapped filtered reads to the *Inga leiocalycina* reference genome ^96^ using the default parameters of *bwa mem2 v2.2*^97^, following which BAM files were sorted and mapping statistics were calculated with *samtools v1.18* ^98^. PCR duplicates were removed from BAM files using *picard MarkDuplicates* v2.26.10 (http://broadinstitute.github.io/picard), and unique molecular identifiers (UMIs) were used to remove further PCR duplicates with *UMI-tools dedup v1.1.4* ^99^. Variants were called per chromosome using the *bcftools* v1.17 ^98^ *mpileup* and *call* commands. We specified that each output VCF file should contain allelic depth, genotype depth and strand bias annotations (*-a AD,DP,SP*), a minimum base quality (*-Q*) and mapping quality (*-q*) of 30, while using a multiallelic caller (*-m*) to retain only variant sites (*-v*) and include annotations of genotype quality (*-f GQ*). Finally, we used *vcftools* v0.1.16 ^100^ to calculate summary statistics for each VCF file, denoting mean depth, base quality, amount of missing data, heterozygosity and allele frequency per site, as well as mean depth and the amount of missing data per individual. These statistics were then plotted using *ggplot2* ^101^ in R v.4.2.1 ^102^.

Each VCF file was then filtered using *vcftools* to remove indels (*--remove-indels*), while removing sites with more than 50% missing data (*--max-missing*), a minimum quality below 30 (--*minQ*), a minimum depth (*--min-meanDP, --minDP*) below 5x and a maximum depth (*--max-meanDP, --maxDP*) above 50x (calculated as 2.5x mean depth to avoid paralogous sites), as well as sites with a minor allele count below 2 (-*-mac*) to remove variants likely resulting from sequencing error. All downstream analyses were performed on the filtered per-chromosome VCF files unless otherwise specified.

### Assessing population structure

We inferred whole-genome phylogenetic trees for all sequenced accessions to obtain a genome-wide summary of phylogenetic relationships, without relying on single regions that may be incongruent. To do so we filtered our VCF files to only include sites with <20% missing data, and then thinned the VCFs to one site per 1500 bases using *vcftools* (*--max-missing* and *--thin*). We then converted the resulting VCFs to phylip files using the *vcf2phylip* script (https://github.com/edgardomortiz/vcf2phylip). We inferred phylogenetic trees using IQ-TREE v2.1.4-beta ^103^, while finding the best substitution model (*-m MFP*), reducing impact of severe model violations (*-bnni*) and with 1000 ultrafast bootstraps (*-B 1000*). We used the same procedure to produce a phylogenetic tree based on the plastomes of all sequenced *Inga* species. Finally, we assessed cyto-nuclear discordance between the resulting phylogenies using the R function *cophylo()* from the *Phytools* package ^104^. All scripts for running IQ-TREE and cophylo analyses, alongside inferred phylogenetic trees, are held in the folder “1_Trees”, available on Dryad (https://doi.org/10.5061/dryad.k6djh9wnb). Finally, we built neighbour net plots using uncorrected P-distances in SPLITSTREE v4.14.6 ^105^ to visualise patterns of shared genetic variation within each regional *Inga* community at the level of regional communities.

To better understand genomic variation across all sequenced *Inga* accessions, as well as within the regional communities defined above, we used principal component analysis (PCA). We first linkage pruned the VCF file containing all accessions with the ‘--indep-pairwise’ tool in PLINK v1.90b6.21 ^106^ using a window size of 50kb, a window step size of 10 and an R^2^ threshold of 0.1, subsequently filtering sites with >20% missing data. This resulted in a BED file with the same sites represented across all accessions to standardise downstream analyses. We used PLINK to run a PCA (*--pca* flag) for all accessions together, and then subset the BED file to run a PCA for each regional community (each of which was filtered to remove sites missing in >5% of individuals). Due to the number of individuals we analysed in each PCA run, we additionally used the Plotly R package ^107^ to plot interactive PCAs for each of our analyses, with points labelled by species and regional community.

We used the same linkage-pruned master BED file to assess population structure for each regional community with ADMIXTURE v1.3.0 ^108^ using default parameters. We determined the best-fit number of populations (K) for each regional community based on the value with the lowest cross-validation error output by ADMIXTURE after 10 iterations, and plotted the ADMIXTURE outputs in R with custom scripts from Joana Meier’s ‘Speciation Genomics’ GitHub (https://github.com/speciationgenomics/scripts/blob/master/plotADMIXTURE.r).

Because the analysis for the “Peru” regional community had over 250 individuals from 18 *Inga* species, we downsampled the Peru BED file to contain only 150,000 variants (as recommended by the ADMIXTURE manual https://dalexander.github.io/admixture/admixture-manual.pdf), and subsequently ran ADMIXTURE for K values between 2-30. For all other regional communities, we ran ADMIXTURE with K values between 2-20.

### Testing genome-wide introgression

We assessed genome-wide signatures of introgression between all species sampled, as well as within regional communities, using Patterson’s D statistic (i.e. the ‘ABBA-BABA’ test; ^109, 110^) and estimated excess allele sharing with the F_4_ ratio ^111^, both implemented in *Dsuite* v04 ^42^. We estimated D and F_4_ statistics for all possible three-taxon combinations using *Dsuite*’s *Dtriosparallel* function (excluding individuals with uncertain taxonomic placement), using the *Zygia, Abarema* and *Albizia* species we sequenced as the outgroup population (n = 9). Branching order information for each taxon trio was included in the analyses using the nuclear genome IQ-tree, trimmed with the R package ape ^112^ to include one accession per species with the least missing data. We assessed the significance of each test using 20 block jackknife resampling runs and corrected *P-*values using Benjamini–Hochberg correction in RSTATIX ^113^. For the accessions from Brazil, we only included those sampled from Amazonia by removing five individuals from our dataset that were sampled from the Atlantic Forest in southeastern Brazil (three *Inga thibaudiana* and two *I. capitata*). We visualized our D-statistic and F_4_ ratio estimates with Ruby scripts available from https://github.com/mmatschiner.

Older hybridisation events can result in correlated F_4_ ratios between related species, and so we controlled for this by estimating the F_branch_ statistic ^114^, and plotted scores onto the single-accession-per-species IQ-TREE with the DSUITE ‘dtools.py’ utility. We then marked Z-scores >5.12, equivalent to a P-value of 0.01 after correction for multiple testing. The population sets file, detailing assignment of accessions to species in the Dsuite analyses, are shown in Extended Data Table S2. All scripts for running Dsuite analyses, alongside raw results files, are held in the folder “2_Dsuite”, available on Dryad.

### Block-wise demographic inference to understand barriers to gene flow

We next aimed to establish what proportion of the genome may act as ‘barriers’ to gene flow within *Inga* syngameons. To do so, we used *gIMble v.1.03* (https://github.com/LohseLab/gIMble; ^46^) to estimate local migration rate (*m_e_*) in genomic windows, while accounting for variation in local effective population size (*N_e_*). We ran *gIMble* on the same species pairs as were used in the *f _dM_* analyses (population selection detailed in Extended Data Table S3), which had elevated gene flow, thus simplifying identification of genomic windows with reduced local migration. We sampled around five co-occurring individuals per species to maximise genome-wide migration rate and reduce the effect of isolation by distance on our inference of barriers to gene flow.

Firstly, we used *gimbleprep* to filter variants from the original VCF file, downsampled to include only the species for each corresponding species-pair analysis. We included only variants at least 2bp away from an indel (to avoid erroneous calls), a minimum quality of 30, a minimum depth of 5x and a maximum depth of twice the average depth (to avoid paralogy issues). Then, we masked repetitive regions in the *Inga leiocalycina* reference genome using RED ^115^, and estimated the genome-wide migration rate for each species pair. To do so we generated blocks of 64 bp (spanning a maximum of 128 bp) across the genome of each species pair. Using these blocks we tested the fit of three global demographic models (strict isolation, migration into species 1 and migration into species 2) for each species pair, and selected the best-fit model for downstream analysis using log composite likelihood (lnCL).

To estimate absolute parameter values for the best-fit demographic model for each species pair, we used a *de-novo* mutation rate for *Inga* of 5x10^−9^ mutat./site/gen. This was inferred using phylogenetic estimates of substitution rate in *Inga* (1 x 10^−3^ subst./site/My) ^30, 89^, which was converted to mutation rate using a generation time of five years based on previous studies of *Inga* demography ^116, 117^. This estimate is congruent with previous work on tropical trees with slower diversification rates (e.g. ^118^) which used mutation rates from *Populus* (2.5 × 10^−9^ mutat./site/gen ^119^). We set the upper parameter bound for divergence time in gIMble (T, measured in generations) based on estimated split times (in years) for each species pair from ^88^, multiplied by 0.2 generations per year, which was then quadrupled to provide a broad upper bound. We used total population size estimates for each *Inga* species (based on plot data from across Amazonia in ^120^), multiplied by 4, to set reasonable upper bounds for effective population size (Ne) in gIMble.

Finally, we estimated variation in local migration rate (*m_e_*) and effective population size (*N_e_*) across the genome for each species pair, using a mean of 21396 blocks, which were a mean of 225799 bases in length. We set the lower bound for *N_e_* to one fifth of the global *N_e_* estimate, and the upper bound to four times the global estimates, following which we conducted the parameter search in 12 grid points with a linear increment. For local *m_e_*, the lower bound was set to zero and the upper bound was set to four times the global estimates, with 16 grid points over the parameter space. We performed this search in *gIMble makegrid*, and then calculated the lnCL for the block-wise site frequency spectrum (bSFS) within windows for each grid-point. The maximum composite likelihood was then recorded for each window, and maximum composite likelihoods were recorded for each window while accounting for each value of local migration (*m_e_*). We then tallied up barriers to gene flow for each species pair by counting windows in which Δ*B* was above zero (Δ*B* = the difference in lnCL between an isolation-with-migration (IM) model with *m_e_* > 0 and a model where m_e_ = 0).

#### Identifying Biosynthetic Gene Clusters in Inga

To ascertain which regions of the genome were associated with chemical defence against herbivores, we annotated the *Inga leiocalycina* reference genome using plantiSMASH (http://plantismash.secondarymetabolites.org/; ^52^) to identify biosynthetic gene clusters (BGC), which are regions of the genome containing co-located genes associated with a certain biosynthetic pathway. We annotated the *I. leiocalycina* genome further using the KEGG Automated Annotation Server (KAAS; https://www.genome.jp/kegg/kaas/; ^121, 122^) to identify genes associated with biosynthesis of flavonoids which are important for chemical defence in *Inga* ^35^. We took this *a-priori* approach using BGCs because the co-located and co-regulated gene clusters held within BGCs have a disproportionate role in secondary metabolism ^51^, which underlies chemical defence that is well established as an important factor governing divergence and ecological coexistence in tropical trees like *Inga* ^7, 36^.

### Estimating introgression and balancing selection in genomic windows

Next, we explored patterns of excess allele sharing, resulting from introgression, across the genome using the *f _dM_* statistic ^53^ in DSUITE. We calculated *f _dM_* for 19 *Inga* species-pairs that showed the most evidence of genome-wide introgression in the D and F_4_ statistics (population selection for *f _dM_* detailed in Extended Data, Table S3). We performed analyses within regional communities for focal species pairs. We calculated *f _dM_* using nonoverlapping windows of 50 informative SNPs with a rolling mean of one window for each focal species pair. Finally, we used custom R scripts to identify outlier windows for each run by extracting *f _dM_* windows below the first percentile of all scores and above the 99^th^ percentile for all scores, as *f _dM_* is centred around zero. All scripts for running fdM analyses and assessing outlying fdM peaks, alongside raw results files, are held in the folder “2_Dsuite”, available on Dryad.

We then assessed evidence of balancing selection across the genome using *BetaScan2* ^123^, which detects clusters of intermediate-frequency alleles surrounding balanced polymorphisms in a sliding window. *BetaScan* assumes no population structure, and so we ran 42 independent *BetaScan* analyses, each consisting of a single *Inga* species in a single regional community (population assignments for each individual used in *BetaScan2* are available in Table S2, Extended Data). Between 5-10 individuals were selected for each run, with population assignments informed by our PCA and ADMIXTURE analyses, within which the individuals with the least missing data were selected. We re-called input VCF files for each *BetaScan* analysis with the same parameters as above but retained invariant sites, and converted each VCF into *BetaScan* format using *glactools* v1.09 ^124^. In each analysis we estimated the standardised beta statistic (BetaSTD) using a window size of 5000bp (based on the average length of a gene in *Inga*) and mutation rate (theta) of 5x10^−9^, based on previous substitution rate estimates in *Inga* ^30,89^. For each analysis we then identified outlying BetaSTD values that exceeded the 99^th^ percentile of all values in R, as above for *f _dM_* scores. For downstream analyses, we standardised genomic windows across all populations for which we estimated balancing selection using *aggregate()* in the R package *dplyr* ^125^. Specifically, we calculated a mean balancing selection score in identical 5kb windows along the genome for each population, which were then Z-score normalised. For candidate regions showing both outliers of *f _dM_*and *BetaSTD*, we plotted scatterplots of these statistics along the genome, highlighting outlying values with different colour points, using *ggplot2* in R.

Finally, we assessed the similarity of balancing selection profiles in chemical defence genes among *Inga* populations. To do so we created two data subsets, first by extracting balancing selection scores from regions of the genome containing PlantiSMASH BGCs, and secondly from regions containing flavonoid genes identified with KEGG only. For each of these subsets, we calculated pairwise Euclidean distance matrices between *Inga* populations using the *dist()* function in *R.* All scripts for running BetaScan2 balancing selection analyses, inference of distance matrixes, plotting and raw results files are held in the folder “3_Balancing_selection_BetaScan2”, available on Dryad.

### Identifying candidate loci for introgression and balancing selection

To understand whether hybridisation generated functional diversity in chemical defence, we identified whether regions of the genome with outlying evidence for introgression and balancing selection contained genes relating to defence chemistry. We annotated outliers with a total of three different annotation types: ‘All genes’ based on the *Inga leiocalycina* reference annotation file, ‘plantiSMASH’ based on the plantiSMASH clusters, alongside ‘KEGG Flavonoid’ based on the KAAS analysis that identified genes associated with flavonoid production. We assigned annotations to outliers based on the distance between the midpoint of a genomic outlier window and the midpoint of each gene or cluster in each annotation type. We took the closest annotated gene for each outlier, retaining only annotations <50 kb from the genomic window midpoint for the ‘All genes’ and ‘KEGG’ annotations, and <2Mb for plantiSMASH clusters (thresholds were based on the average size of a gene or cluster in each annotation type). Finally, we marked genomic outlier windows as occurring within a gene or cluster if its midpoint occurred between the ‘start’ and ‘end’ values of an annotation.

To understand whether specific genomic regions relating to defence chemistry experienced replicated introgression and balancing selection across many species, we used the *geom_histogram()* function in the R package ggplot2 to plot the density of annotated outliers along the genome in 1Mb bins, facet wrapped by chromosome, summing outliers across all analyses for *f _dM_* and betaSTD, alongside. We used the same method to visualise whether outlier frequency was confounded by analytical artefacts such as *f _dM_*window density (since *f _dM_* window size is defined by number of informative SNPs), as well as gene density and repeat density (based on the *Inga leiocalycina* reference genome gff file). All scripts for assessment of overlap between balancing selection scores, introgression outliers and coding genes or BGCs are held in the folder “3_Balancing_selection_BetaScan2”, available on Dryad.

### Assessing introgression of BGC ‘building blocks’

If adaptive introgression generated functional diversity in chemical defence that was then subject to negative density-dependent selection, we would expect introgression of biosynthetic gene cluster (BGC) ‘building blocks’ (as per the Lego Chemistry model of defence evolution), with outlying evidence of both introgression and higher polymorphism as a result of balancing selection.

To explore this, we first identified candidate BGCs by extracting annotated genomic windows that were from the same *Inga* species, the same regional communities, and were within a BGC that had evidence of introgression (*f _dM_* > 0.25) in at least three species pairs. Candidate BGCs also had to be at least >200kb in length, a heuristic value based on the lowest resolution at which we could accurately determine the length of the longest block, and the proportion of the BGC transferred by introgression, based on window size. This resulted in 88 candidate BGC/species pair combinations. To ascertain whether these candidate loci were introgressed and maintained in short linkage blocks, we quantified variation in phylogenetic relationships between species involved in introgression around candidate BGCs using topology weighting in TWISST2 (https://github.com/simonhmartin/twisst2 ^126^). We downsampled the master VCF file to include only individuals and chromosomes identified as candidate BGCs, and created the TWISST2 input file using *sticcs* (https://github.com/simonhmartin/sticcs) with *Zygia pithecoloboides* as outgroup since it had the least missing data. We then inferred local genealogies and their breakpoints along the genome with TWISST2, using a maximum of 100 iterations, and plotted topology weights across each candidate BGC with the ‘plot_twisst.R’ script (https://github.com/simonhmartin/twisst2/blob/main/plot_twisst/plot_twisst.R). We assessed the largest introgressed block length in each BGC by counting the number of concurrent windows with a dominant incongruent genealogy using custom R scripts. We then used box plots in the R package ggplot2 to visualise the longest introgressed block length (both absolute length in kilobases and % of the original BGC introgressed) for all 88 BGC/candidate species trio combinations grouped by BGC type.

Finally, we generated a null expectation of introgression block size for each candidate BGC in each species trio. To do so, we took ten random sequence regions of the same length and chromosome as the candidate BGC for each species trio from our TWISST2 results. We then calculated the maximum introgressed block length for each of the randomly drawn regions with the same method as above. From this, we calculated a mean and standard deviation for the longest block of introgressed variation for each null expectation (i.e., each candidate BGC/species pair combination). All scripts for running TWISST2, calculating the largest introgressed block in a species pair and generation of a null expectation for introgressed block lengths are held in the folder “4_Twisst”, available on Dryad.

### Assessing herbivore turnover and correlation with balancing selection

We aimed to assess whether balancing selection profiles in chemical defence genes were associated with herbivore turnover, as would be expected if defences to which a local herbivore community were naïve experienced negative frequency-dependent selection. To do so, we leveraged herbivore abundance data from our previous work ^22, 76, 127^, collected at the same sites from which the majority of *Inga* samples for genome sequencing were collected for this study. The study sites from which herbivore data were collected were Barro Colorado Island in Panama (9.150°N, 79.850°W), Nouragues research station in French Guiana (4.08°N, 52.683°W) Los Amigos Biological Station in Peru (12.567°S, 70.100W) and Tiputini Biodiversity Station in Ecuador (0.638°S. 76.150°W). In all, these herbivore data represent 16 people-months of field-based data collection per site, and comprise 5,740 individual herbivores belonging to 522 MOTUs (molecular operational taxonomic units) from 13 insect families, collected from 126 *Inga* species.

Herbivore data were collected, DNA barcoded and allocated to MOTUs as described in ^22, 76, 127^. Briefly, in each sampling location, the abundance of all herbivore morphospecies feeding on newly flushing *Inga* leaves was recorded, targeting approximately the same number of flushing leaves per *Inga* species (*ca.* 60). Then, for solitary herbivore species, one individual from each herbivore morphospecies found on a leaf was collected for DNA barcoding. For gregarious herbivore species, at least three individuals per group were collected to ensure that they belonged to the same MOTU. Following collection, a 645 base pair (bp) fragment of the mitochondrial gene cytochrome oxidase I (*COI*) was sequenced for each herbivore collected. DNA barcode sequences from each individual herbivore were allocated to MOTUs using jMOTU v1.0.8 ^128^ and ABGD (Automatic Barcode Gap Discovery ^129^), both of which gave highly concordant MOTU identifications. Because mitochondrial haplotypes can give misleading indications of species membership in sawflies ^130, 131^, multiple sawfly candidate MOTUs were additionally sequenced at two nuclear loci: *wingless* (coding, 327 bp) and *ITS2* (non-coding, 609 bp). The relationships between MOTUs inferred using *COI* data were highly congruent with those based on the two nuclear barcoding loci. MOTUs were subsequently allocated to taxonomic families and/or superfamilies by querying the resulting consensus sequence against the BOLDSYSTEMS database (http://boldsystems.org) using the NCBI BLAST web interface (https://blast.ncbi.nlm.nih.gov/Blast.cgi), with a minimum accepted similarity of 90% for assignment to families. The final, processed herbivore dataset detailed the abundance of each herbivore MOTU collected from all *Inga* host species per regional community (dataset “*Herb_x_com*”), and one that detailed the abundances of each herbivore MOTU found on specific *Inga* species in each regional community (dataset “*Herb_x_Inga_x_com*”). We visualised the abundance of each herbivore family present on each *Inga* species at each site using heat maps and bar plots, implemented in *ggplot2* (for scripts, see “5_BrayCurtis_analyses_herbivore/Herbivore_data_clustering.R” on Dryad).

We used PermANOVA (a one-sided permutation test) with the function *adonis2()*, implemented in the *vegan* R package ^132^, to assess herbivore turnover. Specifically, we aimed to assess the proportion of variation in community composition explained by regional community and *Inga* host species. To do so we performed PermANOVA on our per-*Inga-*species herbivore dataset (dataset “*Herb_x_Inga_x_com*”) and used 100000 permutations for the analysis. Next, we calculated a distance matrix of herbivore similarity for all *Inga* populations using *vegdist()* from the *vegan* R package with Bray-Curtis distances and performed hierarchical clustering with *hclust().* We performed this both for the dataset containing all *Inga* herbivores across regional communities (dataset “*Herb_x _com*”), as well as the per-*Inga*-species dataset (dataset “*Herb_x_Inga_x_com*”). Finally, to test whether turnover in herbivore community per-*Inga* species was correlated with change in balancing selection score in different classes of chemical defence genes (KEGG flavonoid genes and PlantiSMASH BGCs), we performed one-sided Mantel tests ^133^ with the *mantel()* function in R, using 9999 permutations and the Pearson product-moment correlation method. For this, we used the distance matrices derived from balancing selection scores in defence genes, and correlated them with the herbivore turnover distance matrix including the same populations. The raw herbivore data table, describing abundance of herbivore MOTUs from different herbivore families collected on different *Inga*s at different sites, is available on Dryad in the folder “5_BrayCurtis_analyses_herbivore”. Also available in this Dryad folder are scripts for inferring herbivore turnover with Bray-Curtis distances, scripts for PermANOVA analysis and Mantel tests, alongside the resulting Bray-Curtis distance matrices describing herbivore turnover both per site and per site x *Inga* host species combination.

We subsequently aimed to assess phylogenetic turnover in herbivore communities, in order to account for relatedness between herbivore species when estimating herbivore turnover. To do so, we aligned the COI consensus sequence for each MOTU using the program MUSCLE ^134^ for each herbivore family or superfamily (hereafter “family”). We performed alignment and downstream analysis at the family level because of the high chance of homoplasy in short barcode sequences when combining sequence data across families, which could increase the likelihood of incorrect phylogenetic estimation. We generated a total of thirteen COI alignments, comprising all thirteen herbivore families – twelve in the Lepidoptera (Bombycoidea, Gelechioidea, Geometridae, Gracillarioidea, Hesperiidae, Noctuoidea, Pieridae, Pyraloidea, Riodinidae, Tineoidea, Tortricidae and Zygaenoidea) as well as one family in the Hymenoptera (Argidae sawflies). We inferred phylogenetic trees for each COI alignment using IQ-TREE ^103^ by selecting the best-fit substitution model (-MFP) while reducing the impact of severe model violations (-bnni) with 1000 ultrafast bootstrap replicates.

Using each of these family-level trees, and a pared-down herbivore abundance dataset containing only MOTUs from the same family, we estimated the similarity between herbivore communities on each *Inga* species at each site (dataset “*Herb_x_Inga_x_com*”) using the weighted unique fraction metric (UniFrac ^135^) with the *GUniFrac()* function implemented in the R package GUniFrac ^136^. Because UniFrac analyses can be sensitive to uneven herbivore abundances between sites, we removed MOTUs with a total of less than five individuals across all sites. All downstream analyses followed the protocol and parameters outlined above for the herbivore abundance data alone, and each analysis was performed per-herbivore-family. Specifically, we assessed how much variation in phylogenetic community turnover was explained by region and *Inga* host species using PermANOVA, following which we performed hierarchical clustering with *hclust(),* using the UniFrac distances. Then, we used one-sided Mantel tests to assess whether phylogenetic turnover in herbivore community (per-*Inga* species) was correlated with change in balancing selection score in KEGG flavonoid genes and PlantiSMASH BGCs. For the UniFrac PermANOVA tests, we excluded the Bombycoidea, Tineoidea and Zygaenoidea families, which lacked enough data for permutation. For the UniFrac Mantel tests we excluded the same families as for PermANOVA, in addition to the Hesperidae, Gracillarioidea and Pieridae, again due to lack of data. We applied Bonferroni correction ^137^ to all *P-*values for each of our family-level analyses. Included in the folder “6_UNIFRAC_analyses_herbivore_family” on Dryad are scripts for inferring herbivore turnover with UNIFRAC distances, scripts for PermANOVA analysis and Mantel tests, alongside DNA sequence alignments and phylogenetic trees for each herbivore family. Also included are UNIFRAC distance matrices, which describe phylogenetic turnover in herbivores both per site and per site x *Inga* host species combination for 10 herbivore families.

## Data availability

The datasets that support the findings of this study are available from online repositories. All raw reads generated by whole-genome resequencing are available on the European Nucleotide Archive (ENA) under the study accession number PRJEB76593. All sample accession codes are given in Table S1. Other data have been deposited on Dryad (https://doi.org/10.5061/dryad.k6djh9wnb). DNA sequences for the herbivore samples are available from the International Barcode of Life database under the sample IDs IngaHerbiv_0001–IngaHerbiv_2143 and RCMJE_LA01–RCMJE_LA285.

## Code availability

Code underlying the analyses in this paper have been deposited on Dryad (https://doi.org/10.5061/dryad.k6djh9wnb).

## Acknowledgements

We thank the Servicio Nacional Forestal y de Fauna Silvestre (SERFOR, Peru) for granting the research and collection permit in Peru that enabled this study, issued under Directoral Resolution N° RD-000072-2022-DGGSPFFS-DGSPF. We are also thankful to the Brazilian Ministry of Environment for Inga collection and access to genetic resources permits (SISBIO/MMA/Brazil no. 43618-5 and SISGEN/MMA/Brazil no. R7A9E4A). Many thanks to the Ecuadorian Ministerio del Ambiente y Agua for issuing research permissions under the number MAATE DBI-CM-2021-0187. We thank Aniceto Daza Yomona for his fieldwork help in Peru. The authors would also like to acknowledge the Research/Scientific Computing teams at The James Hutton Institute and NIAB for providing computational resources and technical support for the “UK’s Crop Diversity Bioinformatics HPC” (BBSRC grant BB/S019669/1), use of which has contributed to the results reported within this paper. Thanks also to Colin Hughes for his help in the development of the project. We would like to extend our gratitude to Karen Moore, Audrey Farbos, Paul O’Neill, Jemima Onime and the whole team at the University of Exeter Sequencing Service for their help with generating the whole-genome resequencing data for this manuscript. We thank the Wellcome Sanger Institute Tree of Life simple handling, core laboratory, assembly and genome curation teams for production of the reference *Inga* genomes, and the Wellcome Sanger Institute Sequencing Operations long read team for genome sequencing. This manuscript was deeply inspired by the life and work of the late Prof. Thomas A. Kursar.

## Author contributions

Conceptualization: R.J.S., A.D.T., M.-J.E., D.L.F., J.N., G.N.S., F.F.P., P.D.C., C.K., K.G.D., R.T.P.

Methodology: R.J.S., A.D.T., M.-J.E., D.L.F., J.N., G.N.S., M.L., F.F.P., C.H., T.C.M., S.M., J.W., C.Z., P.D.C., C.K., K.G.D., R.T.P.

Investigation: R.J.S., A.D.T., M.-J.E., D.L.F., J.N., G.N.S., M.L., F.F.P., C.H., T.C.M., S.M.,

J.W., C.Z., P.D.C., C.K., K.G.D., R.T.P.

Visualization: R.J.S.

Funding acquisition: R.J.S., A.D.T., G.N.S., M.B., P.D.C., C.K., K.G.D., R.T.P.

Project administration: R.J.S., A.D.T., M.-J.E., A.A.W.S., C.R., G.N.S., M.B., P.D.C., C.K., K.G.D., R.T.P.

Resources: R.J.S., M.-J.E., D.L.F., J.N., G.N.S., M.B., C.H., T.C.M., S.M., J.W., C.Z., H.C.d.L., D.M.N., M.R.L., L.P.d.Q., P.D.C., C.K., K.G.D., R.T.P.

Supervision: A.D.T., M.-J.E., C.R., G.N.S., P.D.C., C.K., K.G.D., R.T.P.

Writing – original draft: R.J.S., A.D.T., J.N., F.F.P., C.K., K.G.D., R.T.P.

Writing – review & editing: R.J.S., A.D.T., M.-J.E., D.L.F., A.A.W.S., C.R., J.N., G.N.S., M.B., M.L., F.F.P., H.C.d.L., D.M.N., M.R.L., L.P.d.Q., P.D.C., C.K., K.G.D., R.T.P.

## Supplementary Information

Supplementary Information is available for this paper.

Available as a single PDF in Extended Data:

• Figs. S1 - S2 and S4 - S11

Available as separate files in Extended Data:

• Tables S1 - S14

• Fig. S3a - S3f and S12 - S14

**Materials & Correspondence**

Correspondence and material requests should be addressed to Rowan J. Schley.

## Main References

1. Koenen, E. J., Clarkson, J. J., Pennington, T. D. & Chatrou, L. W. Recently evolved diversity and convergent radiations of rainforest mahoganies (Meliaceae) shed new light on the origins of rainforest hyperdiversity. New Phytol. 207, 327–339 (2015).

2. Erkens, R. H., Chatrou, L. W., Maas, J. W., van der Niet, T. & Savolainen, V. A rapid diversification of rainforest trees (*Guatteria*; Annonaceae) following dispersal from Central into South America. Mol. Phylogenet. Evol. 44, 399–411 (2007).

3. Dexter, K. G. & Chave, J. Evolutionary patterns of range size, abundance and species richness in Amazonian angiosperm trees. PeerJ 4, e2402 (2016).

4. Schluter, D. Evidence for ecological speciation and its alternative. Science 323, 737–741 (2009).

5. Wellborn, G. A. & Langerhans, R. B. Ecological opportunity and the adaptive diversification of lineages. Ecology and evolution 5, 176–195 (2015).

6. Yoder, J. B. et al. Ecological opportunity and the origin of adaptive radiations. J. Evol. Biol. 23, 1581–1596 (2010).

7. Forrister, D. L. et al. Diversity and divergence: evolution of secondary metabolism in the tropical tree genus *Inga*. New Phytol. 237, 631–642 (2023).

8. Pennington, R. T., Hughes, M. & Moonlight, P. W. The origins of tropical rainforest hyperdiversity. Trends Plant Sci. 20, 693–695 (2015).

9. Hughes, C. E., Nyffeler, R. & Linder, H. P. Evolutionary plant radiations: where, when, why and how? New Phytol. 207, 249–253 (2015).

10. Ulloa Ulloa, C. et al. An integrated assessment of the vascular plant species of the Americas. Science 358, 1614–1617 (2017).

11. Bass, M. S. et al. Global conservation significance of Ecuador’s Yasuní National Park. PloS one 5, e8767 (2010).

12. Valencia, R., Balslev, H. & Miño, G. P. Y. High tree alpha-diversity in Amazonian Ecuador. Biodiversity & Conservation 3, 21–28 (1994).

13. Rivers, M. et al. European Red List of Trees. (2019).

14. Guevara-Andino, J. E. et al. Neotropics as a cradle for adaptive radiations. Cold Spring Harbor Perspectives in Biology 17, a041452 (2025).

15. Bagchi, R. et al. Pathogens and insect herbivores drive rainforest plant diversity and composition. Nature 506, 85 (2014).

16. Janzen, D. H. Herbivores and the number of tree species in tropical forests. Am. Nat. 104, 501–528 (1970).

17. Connell, J. H. in Dynamics of Populations (eds Den Boer, P. J. & Gradwell, G. R.) (Centre for Agricultural Publishing and Documentation, Wageningen, The Netherlands, 1971).

18. Forrister, D. L., Endara, M., Younkin, G. C., Coley, P. D. & Kursar, T. A. Herbivores as drivers of negative density dependence in tropical forest saplings. Science 363, 1213–1216 (2019).

19. Fine, P. V., Mesones, I. & Coley, P. D. Herbivores promote habitat specialization by trees in Amazonian forests. Science 305, 663–665 (2004).

20. Sedio, B. E., Rojas Echeverri, J. C., Boya P, C. A. & Wright, S. J. Sources of variation in foliar secondary chemistry in a tropical forest tree community. Ecology 98, 616–623 (2017).

21. Bialic-Murphy, L. et al. The pace of life for forest trees. Science 386, 92–98 (2024).

22. Endara, M. et al. Coevolutionary arms race versus host defense chase in a tropical herbivore–plant system. Proceedings of the National Academy of Sciences 114, E7499–E7505 (2017).

23. Buck, R. & Flores-Rentería, L. The syngameon enigma. Plants 11, 895 (2022).

24. Cervantes, S., Schley, R., Hardy, O. J. & Ojeda, D. Do syngameons exist in tropical trees? Challenges to determine their existence and estimate their frequency. Biology Letters 21, 20250444 (2025).

25. Meier, J. I. et al. Cycles of fusion and fission enabled rapid parallel adaptive radiations in African cichlids. Science 381, eade2833 (2023).

26. Cannon, C. H. & Petit, R. J. The oak syngameon: more than the sum of its parts. New Phytol. 226, 978–983 (2020).

27. Ashton, P. S. Speciation among tropical forest trees: some deductions in the light of recent evidence. Biol. J. Linn. Soc. 1, 155–196 (1969).

28. Abbott, R. J. Plant speciation across environmental gradients and the occurrence and nature of hybrid zones. J Syst Evol 55, 238–258 (2017).

29. Schley, R. J., Twyford, A. D. & Pennington, R. T. Hybridization: a ‘double-edged sword’ for Neotropical plant diversity. Bot. J. Linn. Soc. 199, 331–356 (2022).

30. Richardson, J. E., Pennington, R. T., Pennington, T. D. & Hollingsworth, P. M. Rapid diversification of a species-rich genus of neotropical rain forest trees. Science 293, 2242–2245 (2001).

31. Pennington, T. D. in The Genus Inga: Botany. (Royal Botanic Gardens, 1997).

32. Ringelberg, J. J. et al. Precipitation is the main axis of tropical plant phylogenetic turnover across space and time. Science Advances 9, eade4954 (2023).

33. Valencia, R. et al. in Tropical forest diversity and dynamism: findings from a large-scale plot network (eds Losos, E. & Leigh, E. G. L.) 620 (University of Chicago Press Chicago, 2004).

34. Endara, M. et al. Chemocoding as an identification tool where morphological and DNA based methods fall short: Inga as a case study. New Phytol. 218, 847–858 (2018).

35. Coley, P. D., Endara, M. & Kursar, T. A. Consequences of interspecific variation in defenses and herbivore host choice for the ecology and evolution of *Inga*, a speciose rainforest tree. Oecologia 187, 361–376 (2018).

36. Kursar, T. A. et al. The evolution of antiherbivore defenses and their contribution to species coexistence in the tropical tree genus *Inga*. Proc. Natl. Acad. Sci. U. S. A. 106, 18073–18078 (2009).

37. Schley, R. J. et al. Rampant Reticulation in a Rapid Radiation of Tropical Trees - Insights from *Inga* (Fabaceae). Syst. Biol. (2025).

38. Rollo, A. et al. Genetic diversity and hybridization in the two species *Inga ingoides* and *Inga edulis*: potential applications for agroforestry in the Peruvian Amazon. Ann. For. Sci. 73, 425–435 (2016).

39. Koptur, S. Flowering phenology and floral biology of *Inga* (Fabaceae: Mimosoideae). Syst. Bot., 354–368 (1983).

40. Donoghue, M. J. & Edwards, E. J. Model clades are vital for comparative biology, and ascertainment bias is not a problem in practice: a response to Beaulieu and O’Meara (2018). Am. J. Bot. 106, 327–330 (2019).

41. Marques, D. A., Meier, J. I. & Seehausen, O. A combinatorial view on speciation and adaptive radiation. Trends in Ecology & Evolution 34, 531–544 (2019).

42. Malinsky, M., Matschiner, M. & Svardal, H. Dsuite fast D statistics and related admixture evidence from VCF files. Molecular Ecology Resources 21, 584–595 (2021).

43. Buck, R. et al. Sequential hybridization may have facilitated ecological transitions in the Southwestern pinyon pine syngameon. New Phytol. 237, 2435–2449 (2023).

44. Rieseberg, L. H. & Soltis, D. E. Phylogenetic consequences of cytoplasmic gene flow in plants. Evolutionary Trends in Plants 5, 65–84 (1991).

45. Seehausen, O. Hybridization and adaptive radiation. Trends in Ecology & Evolution 19, 198–207 (2004).

46. Laetsch, D. R. et al. Demographically explicit scans for barriers to gene flow using gIMble. PLoS genetics 19, e1010999 (2023).

47. Ma, T. et al. Ancient polymorphisms and divergence hitchhiking contribute to genomic islands of divergence within a poplar species complex. Proceedings of the National Academy of Sciences 115, E236–E243 (2018).

48. Mackintosh, A. et al. Chromosome fissions and fusions act as barriers to gene flow between *Brenthis* fritillary butterflies. Mol. Biol. Evol. 40, msad043 (2023).

49. Wogan, G. O., Yuan, M. L., Mahler, D. L. & Wang, I. J. Hybridization and transgressive evolution generate diversity in an adaptive radiation of Anolis lizards. Syst. Biol. 72, 874–884 (2023).

50. Sherman, D. H. The Lego-ization of polyketide biosynthesis. Nat. Biotechnol. 23, 1083–1084 (2005).

51. Cawood, G. L. & Ton, J. Decoding resilience: ecology, regulation, and evolution of biosynthetic gene clusters. Trends Plant Sci. 30, 185–198 (2025).

52. Kautsar, S. A., Suarez Duran, H. G., Blin, K., Osbourn, A. & Medema, M. H. plantiSMASH: automated identification, annotation and expression analysis of plant biosynthetic gene clusters. Nucleic Acids Res. 45, W55–W63 (2017).

53. Malinsky, M. et al. Genomic islands of speciation separate cichlid ecomorphs in an East African crater lake. Science 350, 1493–1498 (2015).

54. Kawai, Y., Ono, E. & Mizutani, M. Evolution and diversity of the 2– oxoglutarate dependent dioxygenase superfamily in plants. The Plant Journal 78, 328–343 (2014).

55. Peng, M. et al. Differentially evolved glucosyltransferases determine natural variation of rice flavone accumulation and UV-tolerance. Nature communications 8, 1975 (2017).

56. Wei, S., Zhang, W., Fu, R. & Zhang, Y. Genome-wide characterization of 2-oxoglutarate and Fe (II)-dependent dioxygenase family genes in tomato during growth cycle and their roles in metabolism. BMC Genomics 22, 126 (2021).

57. Breiteneder, H. & Kraft, D. The history and science of the major birch pollen allergen Bet v 1. Biomolecules 13, 1151 (2023).

58. Muro-Villanueva, F. & Nett, R. S. Emerging functions within the enzyme families of plant alkaloid biosynthesis. Phytochemistry Reviews, 1–22 (2023).

59. Zhou, F. & Pichersky, E. The complete functional characterisation of the terpene synthase family in tomato. New Phytol. 226, 1341–1360 (2020).

60. Maeda, H. & Dudareva, N. The shikimate pathway and aromatic amino acid biosynthesis in plants. Annual review of plant biology 63, 73–105 (2012).

61. Edelman, N. B. et al. Genomic architecture and introgression shape a butterfly radiation. Science 366, 594–599 (2019).

62. Schumer, M. et al. Natural selection interacts with recombination to shape the evolution of hybrid genomes. Science 360, 656–660 (2018).

63. Nützmann, H., Huang, A. & Osbourn, A. Plant metabolic clusters–from genetics to genomics. New Phytol. 211, 771–789 (2016).

64. Guo, L. et al. The opium poppy genome and morphinan production. Science 362, 343–347 (2018).

65. Winzer, T. et al. A *Papaver somniferum* 10-gene cluster for synthesis of the anticancer alkaloid noscapine. Science 336, 1704–1708 (2012).

66. Singh, P. et al. Highly modular genomic architecture underlies combinatorial mechanism of speciation and adaptive radiation. bioRxiv, 2025.07. 07.663194 (2025).

67. Connell, J. H. Diversity in tropical rain forests and coral reefs: high diversity of trees and corals is maintained only in a nonequilibrium state. Science 199, 1302–1310 (1978).

68. Suarez-Gonzalez, A., Lexer, C. & Cronk, Q. C. B. Adaptive introgression: a plant perspective. Biol. Lett. 14, 10.1098/rsbl.2017.0688 (2018).

69. Delph, L. F. & Kelly, J. K. On the importance of balancing selection in plants. New Phytol. 201, 45–56 (2014).

70. The Heliconius Genome Consortium. Butterfly genome reveals promiscuous exchange of mimicry adaptations among species. Nature 487, 94–98 (2012).

71. Schierup, M. H., Vekemans, X. & Charlesworth, D. The effect of subdivision on variation at multi-allelic loci under balancing selection. Genetics Research 76, 51–62 (2000).

72. Castric, V., Bechsgaard, J., Schierup, M. H. & Vekemans, X. Repeated adaptive introgression at a gene under multiallelic balancing selection. PLoS Genetics 4, e1000168 (2008).

73. Whitney, K. D., Randell, R. A. & Rieseberg, L. H. Adaptive introgression of herbivore resistance traits in the weedy sunflower *Helianthus annuus*. Am. Nat. 167, 794–807 (2006).

74. Barraclough, T. G. Does selection favour the maintenance of porous species boundaries? J. Evol. Biol. 37, 616–627 (2024).

75. Becerra, J. X. The impact of herbivore–plant coevolution on plant community structure. Proceedings of the National Academy of Sciences 104, 7483–7488 (2007).

76. Endara, M. et al. Tracking of host defenses and phylogeny during the radiation of neotropical *Inga*-feeding sawflies (Hymenoptera; Argidae). Frontiers in Plant Science 9, 1237 (2018).

77. Carley, L. N. et al. Ecological factors influence balancing selection on leaf chemical profiles of a wildflower. Nature ecology & evolution 5, 1135–1144 (2021).

78. Gloss, A. D., Dittrich, A. C. N., Goldman-Huertas, B. & Whiteman, N. K. Maintenance of genetic diversity through plant–herbivore interactions. Curr. Opin. Plant Biol. 16, 443–450 (2013).

79. Dexter, K. G. et al. Dispersal assembly of rain forest tree communities across the Amazon basin. Proc. Natl. Acad. Sci. U. S. A. 114, 2645–2650 (2017).

80. Endara, M. et al. The role of plant secondary metabolites in shaping regional and local plant community assembly. J. Ecol. 110, 34–45 (2022).

81. Ehrlich, P. R. & Raven, P. H. Butterflies and plants: a study in coevolution. Evolution, 586–608 (1964).

82. Fine, P. V. et al. Insect herbivores, chemical innovation, and the evolution of habitat specialization in Amazonian trees. Ecology 94, 1764–1775 (2013).

83. War, A. R. et al. Effect of plant secondary metabolites on legume pod borer, *Helicoverpa armigera*. Journal of Pest Science 86, 399–408 (2013).

84. Barrier, M., Baldwin, B. G., Robichaux, R. H. & Purugganan, M. D. Interspecific hybrid ancestry of a plant adaptive radiation: allopolyploidy of the Hawaiian silversword alliance (Asteraceae) inferred from floral homeotic gene duplications. Mol. Biol. Evol. 16, 1105–1113 (1999).

85. Lamichhaney, S. et al. Rapid hybrid speciation in Darwin’s finches. Science 359, 224–228 (2018).

86. Koptur, S. Outcrossing and pollinator limitation of fruit set: breeding systems of neotropical *Inga* trees (Fabaceae: Mimosoideae). Evolution 38, 1130–1143 (1984).

## Methods references

87. WCVP. World Checklist of Vascular Plants, version 2.0. Facilitated by the Royal Botanic Gardens, Kew. Published on the Internet; http://wcvp.science.kew.org/ *Retrieved 31st January 2020*. <https://Published on the Internet; http://wcvp.science.kew.org/>(2020).

88. Nicholls, J. A. et al. Continuous colonization of the Atlantic coastal rain forests of South America from Amazônia. Proceedings of the Royal Society B. 291.

89. Nicholls, J. A. et al. Using targeted enrichment of nuclear genes to increase phylogenetic resolution in the neotropical rain forest genus Inga (Leguminosae: Mimosoideae). Frontiers in Plant Science 6, 710 (2015).

90. Schley, R. J. et al. The frequency and importance of polyploidy in tropical rainforest tree radiations. New Phytol. (2025).

91. Hanson, L. Some new chromosome counts in the genus *Inga* (Leguminosae: Mimosoideae). Kew Bull., 801–804 (1995).

92. Figueiredo, M. F. et al. Intraspecific and interspecific polyploidy of Brazilian species of the genus *Inga* (Leguminosae: Mimosoideae). Genetics and Molecular Research 13, 3395–3403 (2014).

93. Chen, S. Ultrafast one pass FASTQ data preprocessing, quality control, and deduplication using fastp. Imeta 2, e107 (2023).

94. Andrews, S. FastQC: a quality control tool for high throughput sequence data. Available online at http://www.bioinformatics.babraham.ac.uk/projects/fastqc. 0.11.9 (2010).

95. Ewels, P., Magnusson, M., Lundin, S. & Käller, M. MultiQC: summarize analysis results for multiple tools and samples in a single report. Bioinformatics 32, 3047–3048 (2016).

96. Schley, R. J. et al. The genome sequence of *Inga leiocalycina* Benth. Wellcome Open Research 9, 606 (2024).

97. Vasimuddin, M., Misra, S., Li, H. & Aluru, S. Efficient architecture-aware acceleration of BWA-MEM for multicore systems (2019 IEEE international parallel and distributed processing symposium (IPDPS), IEEE, 2019).

98. Danecek, P. et al. Twelve years of SAMtools and BCFtools. Gigascience 10, giab008 (2021).

99. Smith, T., Heger, A. & Sudbery, I. UMI-tools: modeling sequencing errors in Unique Molecular Identifiers to improve quantification accuracy. Genome Res. 27, 491–499 (2017).

100. Danecek, P. et al. The variant call format and VCFtools. Bioinformatics 27, 2156–2158 (2011).

101. Wickham, H. ggplot2: elegant graphics for data analysis. 3.3.6 (2016).

102. R Development Core Team. R: A language and environment for statistical computing. Available at: http://www.R-project.org/. **3.6** (2013).

103. Nguyen, L., Schmidt, H. A., Von Haeseler, A. & Minh, B. Q. IQ-TREE: a fast and effective stochastic algorithm for estimating maximum-likelihood phylogenies. Mol. Biol. Evol. 32, 268–274 (2015).

104. Revell, L. J. phytools: an R package for phylogenetic comparative biology (and other things). Methods in Ecology and Evolution 3, 217–223 (2012).

105. Huson, D. H. & Bryant, D. Application of phylogenetic networks in evolutionary studies. Mol. Biol. Evol. 23, 254–267 (2005).

106. Purcell, S. et al. PLINK: a tool set for whole-genome association and population-based linkage analyses. The American journal of human genetics 81, 559–575 (2007).

107. Plotly Technologies Inc. Collaborative data science. (2015).

108. Alexander, D. H., Novembre, J. & Lange, K. Fast model-based estimation of ancestry in unrelated individuals. Genome Res. 19, 1655–1664 (2009).

109. Green, R. E. et al. A draft sequence of the Neandertal genome. Science 328, 710–722 (2010).

110. Durand, E. Y., Patterson, N., Reich, D. & Slatkin, M. Testing for ancient admixture between closely related populations. Mol. Biol. Evol. 28, 2239–2252 (2011).

111. Patterson, N. et al. Ancient admixture in human history. Genetics 192, 1065–1093 (2012).

112. Paradis, E. & Schliep, K. ape 5.0: an environment for modern phylogenetics and evolutionary analyses in R. Bioinformatics 35, 526–528 (2019).

113. Kassambara, A. rstatix: Pipe-friendly framework for basic statistical tests. CRAN: Contributed Packages (2019).

114. Malinsky, M. et al. Whole-genome sequences of Malawi cichlids reveal multiple radiations interconnected by gene flow. Nature Ecology & Evolution 2, 1940–1955 (2018).

115. Girgis, H. Z. Red: an intelligent, rapid, accurate tool for detecting repeats de-novo on the genomic scale. BMC Bioinformatics 16, 227 (2015).

116. Baker, T. R. et al. Fast demographic traits promote high diversification rates of Amazonian trees. Ecol. Lett. 17, 527–536 (2014).

117. Lojka, B., Preininger, D., Lojkova, J., Banout, J. & Polesny, Z. Biomass growth and farmer knowledge of *Inga edulis* in Peruvian Amazon. Agricultura Tropica y Subtropica 38, 44–51 (2005).

118. Piñeiro, R. et al. Contrasting genetic signal of recolonization after rainforest fragmentation in African trees with different dispersal abilities. Proceedings of the National Academy of Sciences 118, e2013979118 (2021).

119. Ingvarsson, P. K. Multilocus patterns of nucleotide polymorphism 1097 and the demographic history of *Populus tremula*. Genetics 180, 329–340 (2008).

120. ter Steege, H., et al. Hyperdominance in the Amazonian tree flora. Science 342, 1243092 (2013).

121. Moriya, Y., Itoh, M., Okuda, S., Yoshizawa, A. C. & Kanehisa, M. KAAS: an automatic genome annotation and pathway reconstruction server. Nucleic Acids Res. 35, W182–W185 (2007).

122. Kanehisa, M., Furumichi, M., Tanabe, M., Sato, Y. & Morishima, K. KEGG: new perspectives on genomes, pathways, diseases and drugs. Nucleic Acids Res. 45, D353–D361 (2017).

123. Siewert, K. M. & Voight, B. F. BetaScan2: Standardized statistics to detect balancing selection utilizing substitution data. Genome Biology and Evolution 12, 3873–3877 (2020).

124. Renaud, G. glactools: a command-line toolset for the management of genotype likelihoods and allele counts. Bioinformatics 34, 1398–1400 (2018).

125. Wickham, H., François R., Henry, L., Müller, K. & Vaughan D. dplyr: A Grammar of Data Manipulation. *R package version 04*., p156 (2025).

126. Martin, S. H. A model-free method for genealogical inference without phasing and its application for topology weighting. Genetics 232, iyaf181 (2026).

127. Coley, P. D., et al. Macroevolutionary patterns in overexpression of tyrosine: An anti׈herbivore defence in a speciose tropical tree genus, *Inga* (Fabaceae). J. Ecol. 107, 1620–1632 (2019).

128. Jones, M., Ghoorah, A. & Blaxter, M. jMOTU and taxonerator: turning DNA barcode sequences into annotated operational taxonomic units. PLoS one 6, e19259 (2011).

129. Puillandre, N., Lambert, A., Brouillet, S. & Achaz, G. ABGD, Automatic Barcode Gap Discovery for primary species delimitation. Mol. Ecol. 21, 1864–1877 (2012).

130. Prous, M., Heidemaa, M. & Soon, V. *Empria longicornis* species group: taxonomic revision with notes on phylogeny and ecology (Hymenoptera, Tenthredinidae). Zootaxa 2756, 1–39 (2011).

131. Schmidt, S., et al. Identification of sawflies and horntails (Hymenoptera, ‘Symphyta’) through DNA barcodes: successes and caveats. Molecular Ecology Resources 17, 670–685 (2017).

132. Oksanen, J., et al. vegan: Community Ecology Package. R package version 2.5–6. *vegan: Community Ecology Package* 2.5–6 (2019).

133. Mantel, N. The detection of disease clustering and a generalized 1130 regression approach. Cancer Res. 27, 209–220 (1967).

134. Edgar, R. C. MUSCLE: multiple sequence alignment with high accuracy and high throughput. Nucleic Acids Res. 32, 1792–1797 (2004).

135. Lozupone, C. & Knight, R. UniFrac: a new phylogenetic method for comparing microbial communities. Appl. Environ. Microbiol. 71, 8228–8235 (2005).

136. Chen, J., et al. Associating microbiome composition with environmental covariates using generalized UniFrac distances. Bioinformatics 28, 2106–2113 (2012).

137. Bonferroni, C. Teoria statistica delle classi e calcolo delle probabilita. Pubblicazioni del R Istituto Superiore di Scienze Economiche e Commericiali di Firenze 8, 3–62 (1936).

