## Extended Data for "Hybridisation and herbivory fuel Amazonian tree radiations"

**Captions of** [**Extended Data Tables in separate files**	23](#_Toc228530053)

Extended Data Figures

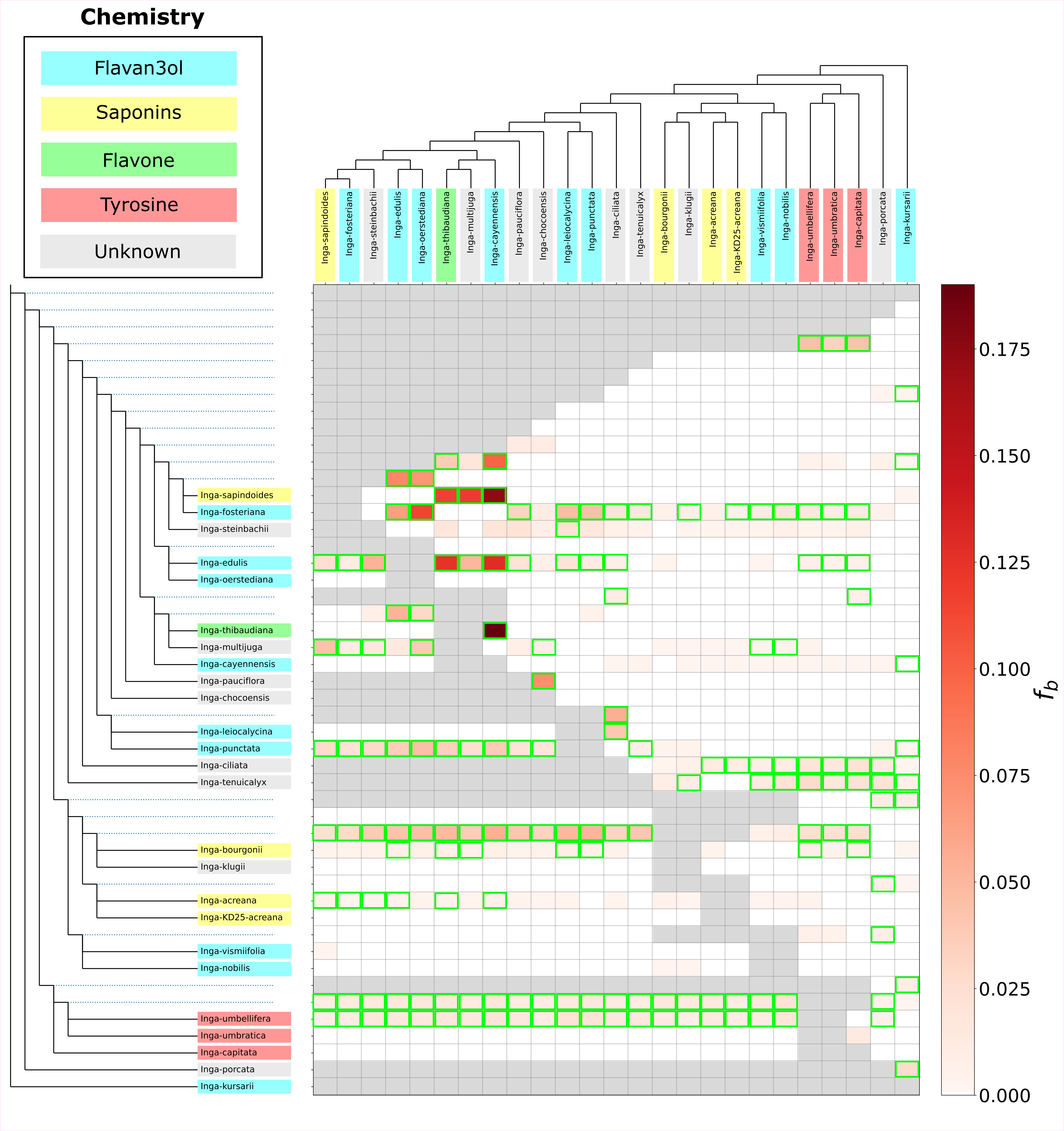

### Figure S1a. Heatmap of ‘Fbranch’ scores plotted for all sequenced Inga species, with all regional communities combined per species.

The colour of each square in the heat map signifies the amount of excess allele sharing (fb; dark red = high estimate). The tree is displayed in an ‘expanded’ form along the y axis, so that all branches (including internal ones, marked as dotted lines), correspond to a row in the heatmap. A summarised version of the nuclear phylogenetic tree from Fig. S5, with one tip per species, is represented on the x axis such that each tip corresponds to a column in the heatmap. Fbranch values with a Z-score >5.12 (corresponding to P<0.01 after multiple-testing correction) are highlighted with a green box. The main mode of chemical defence (where known) used by each species is displayed as a coloured box around the taxon name, after (18). Grey boxes indicate Fbranch tests that were not possible to perform due to the topology of the tree, e.g. between sister species.

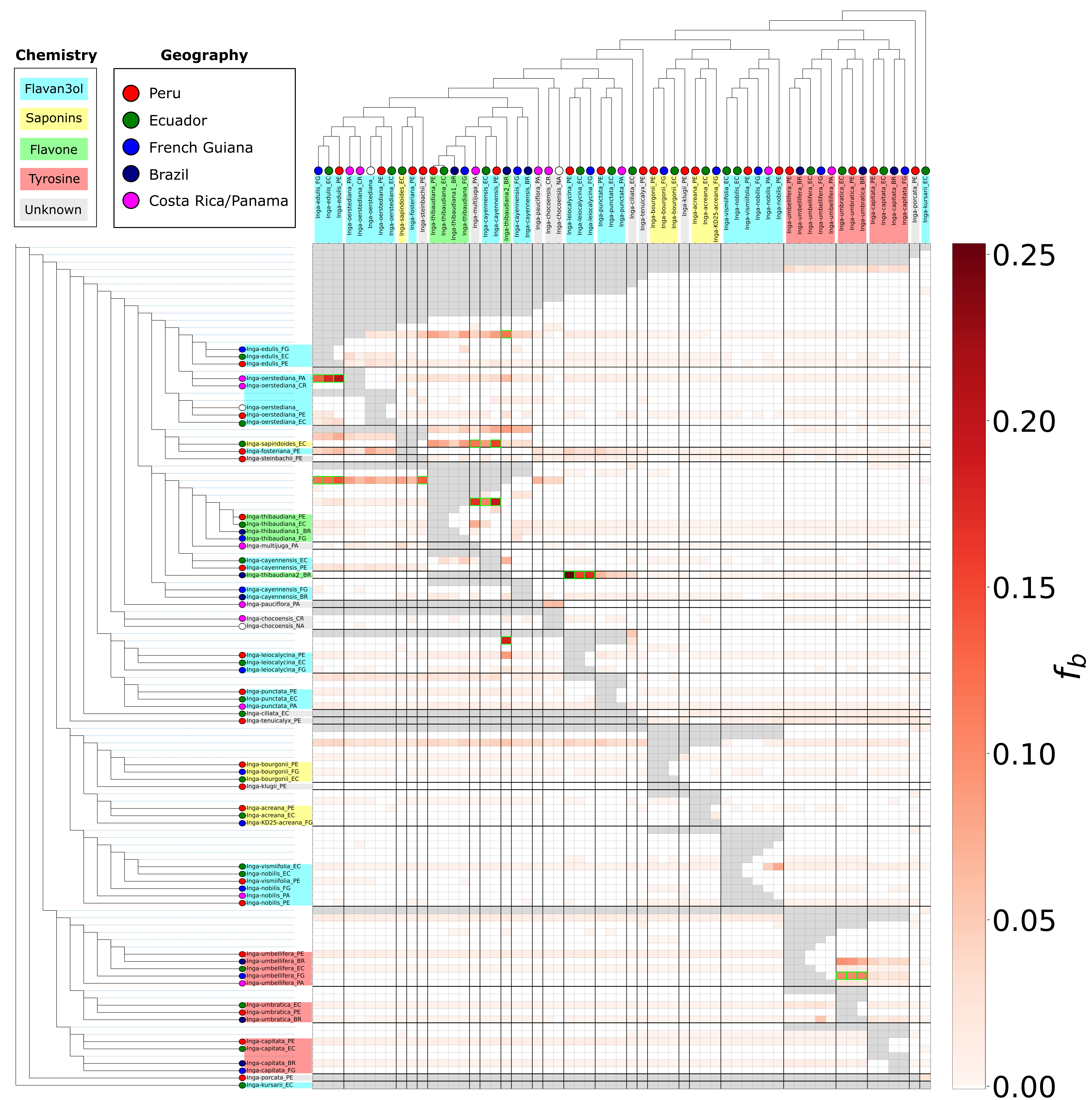

### Figure S1b. Heatmap of ‘Fbranch’ scores plotted for all sequenced Inga species, with all regional communities separated per species.

The colour of each square in the heat map signifies the amount of excess allele sharing (f_b_; dark red = high estimate). The tree is displayed in an ‘expanded’ form along the y axis, so that all branches (including internal ones, marked as dotted lines), correspond to a row in the heatmap. A summarised version of the nuclear phylogenetic tree from Figure S5, with one tip per species, is represented on the x axis such that each tip corresponds to a column in the heatmap. F_branch_ values with a Z-score >5.12 (corresponding to P<0.01 after multiple-testing correction) are highlighted with a green box. The main mode of chemical defence (where known) used by each species is displayed as a coloured box around the taxon name, after *(18)*. Grey boxes indicate F_branch_ tests that were not possible to perform due to the topology of the tree, e.g. between sister species, and within this analysis intra-specific F scores were set to 0 to better visualise introgression.

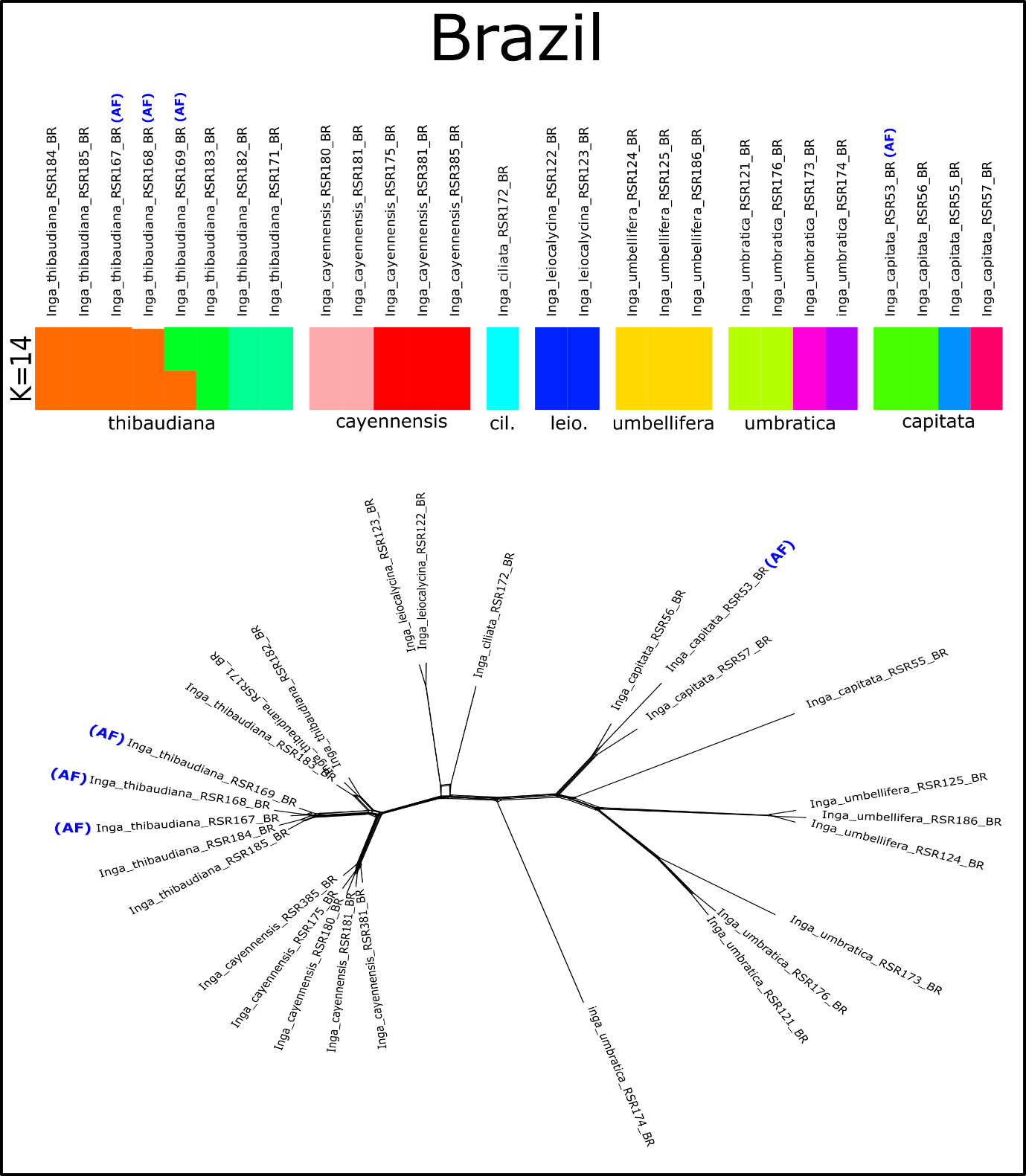

Figure S2a. ADMIXTURE and SPLITSTREE population structure plots for Brazilian samples.
The top panel shows ADMIXTURE plot generated using the K value (i.e., the number of estimated genetic clusters) with the lowest cross-validation error. The ADMIXTURE plot indicates ancestry proportions from K inferred genetic groups, which are indicated with different colours. Each column represents one accession, with accession names above the column. Morphological species identifications for groups of accessions are labelled below the x axis. The bottom panel shows a SplitsTree built using uncorrelated P distances, where connecting edges indicate shared variation. Both plots were inferred using whole-genome resequencing data for Inga accessions collected from Brazil. Accessions collected from the Atlantic Forest in southeastern Brazil are marked with (AF). In the ADMIXTURE plot labels, cil. = *I. ciliata*; leio. = *I. leiocalycina*.

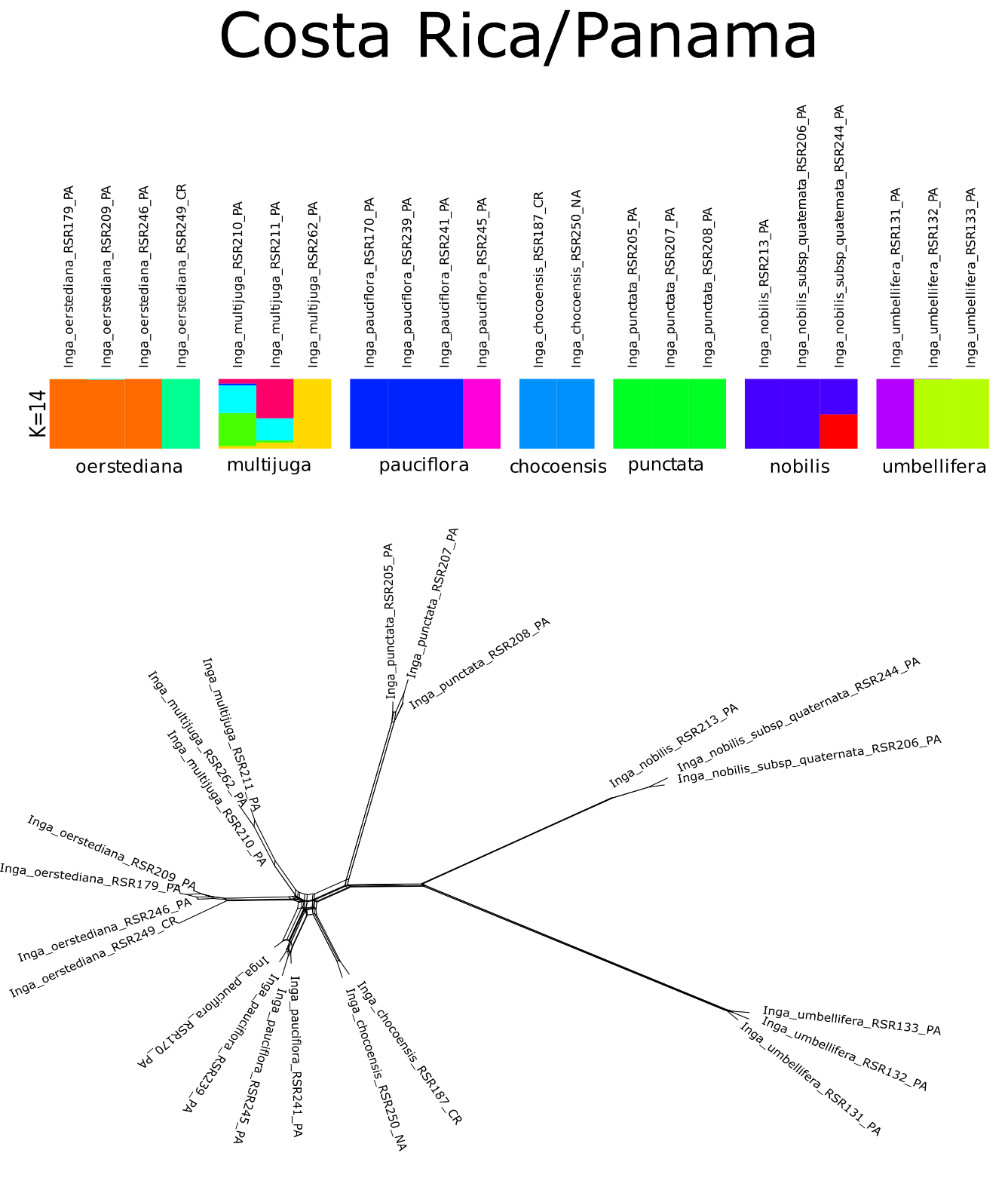

Figure S2b. ADMIXTURE and SPLITSTREE population structure plots for Costa Rican and Panamanian samples.
The top panel shows ADMIXTURE plot generated using the K value (i.e., the number of estimated genetic clusters) with the lowest cross-validation error. The ADMIXTURE plot indicates ancestry proportions from K inferred genetic groups, which are indicated with different colours. Each column represents one accession, with accession names above the column. Morphological species identifications for groups of accessions are labelled below the x axis. The bottom panel shows a SplitsTree built using uncorrelated P distances, where connecting edges indicate shared variation. Both plots were inferred using whole-genome resequencing data for Inga accessions collected from Costa Rica and Panama.

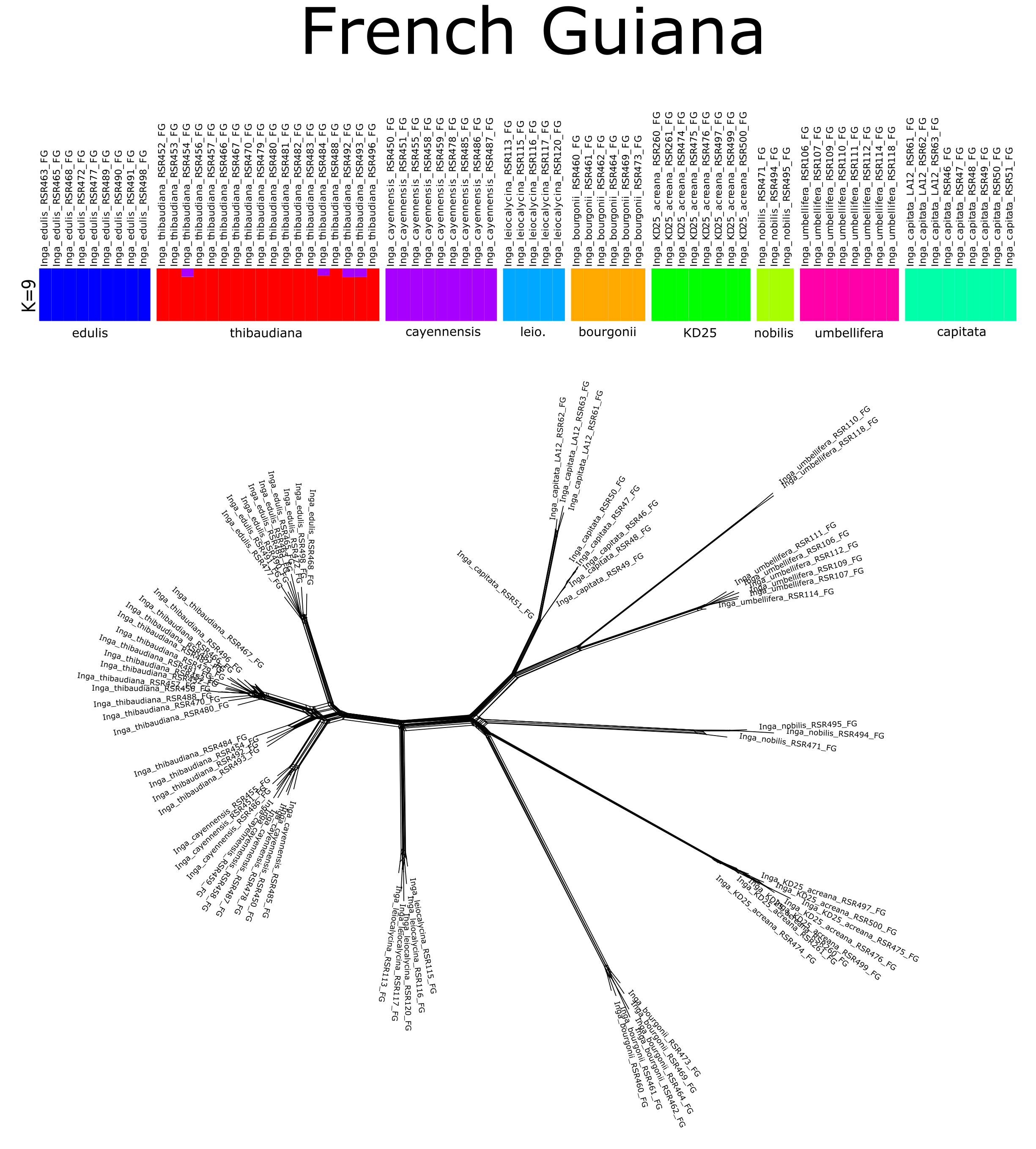
Figure S2c. ADMIXTURE and SPLITSTREE population structure plots for French Guianan samples.
The top panel shows ADMIXTURE plot generated using the K value (i.e., the number of estimated genetic clusters) with the lowest cross-validation error. The ADMIXTURE plot indicates ancestry proportions from K inferred genetic groups, which are indicated with different colours. Each column represents one accession, with accession names above the column. Morphological species identifications for groups of accessions are labelled below the x axis. The bottom panel shows a SplitsTree built using uncorrelated P distances, where connecting edges indicate shared variation. Both plots were inferred using whole-genome resequencing data for Inga accessions collected from French Guiana. In the ADMIXTURE plot labels leio. = *I. leiocalycina*

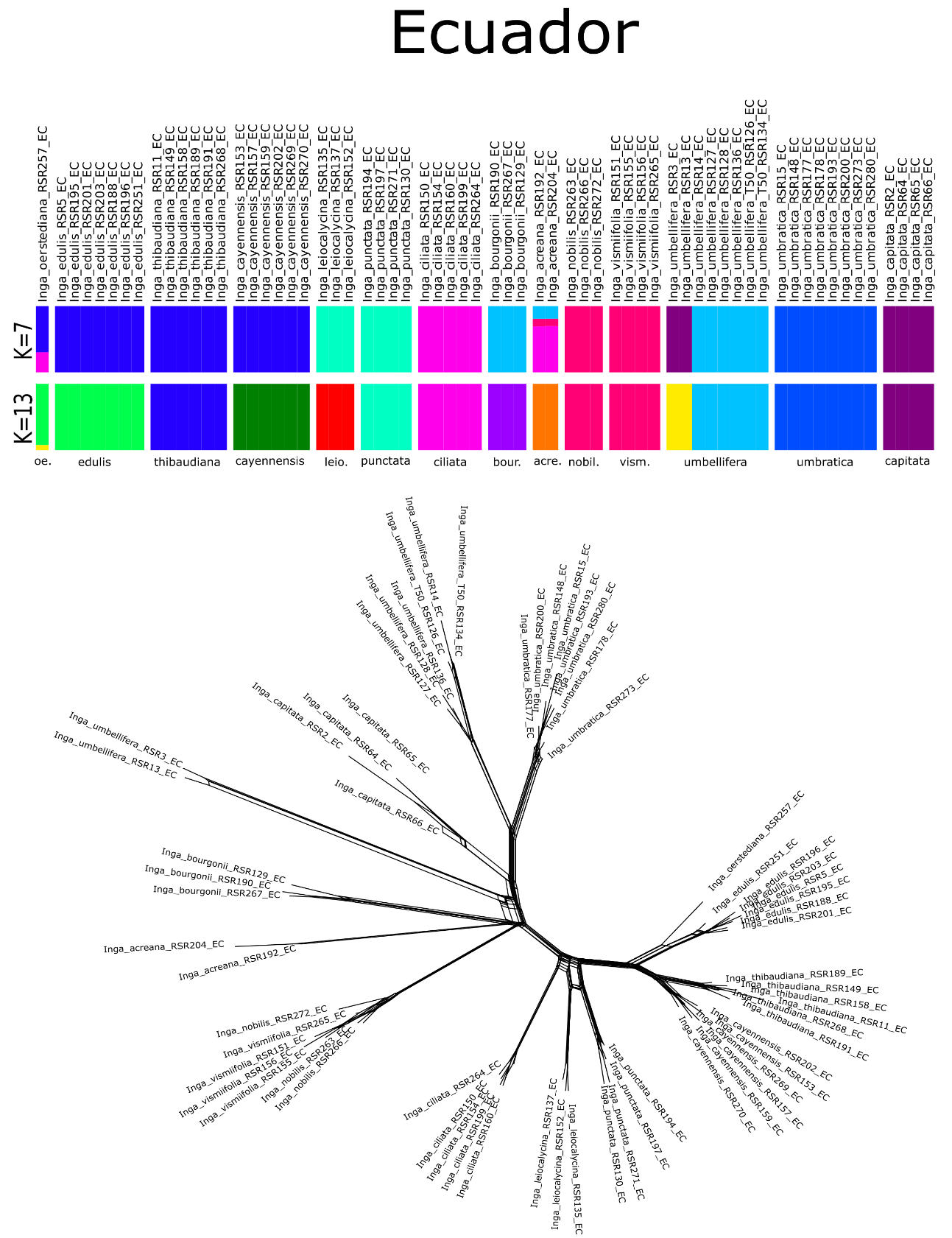

Figure S2d. ADMIXTURE and SPLITSTREE population structure plots for Ecuadorian samples.
The top panel shows ADMIXTURE plot generated using the K values (i.e., the number of estimated genetic clusters) with the lowest cross-validation errors. The ADMIXTURE plot indicates ancestry proportions from K inferred genetic groups, which are indicated with different colours. Each column represents one accession, with accession names above the column. Morphological species identifications for groups of accessions are labelled below the x axis. The bottom panel shows a SplitsTree built using uncorrelated P distances, where connecting edges indicate shared variation. Both plots were inferred using whole-genome resequencing data for Inga accessions collected from Ecuador. In the ADMIXTURE plot labels, oe. = *I. oerstediana*; leio. = *I. leiocalycina*; bour. = *I. bourgonii*; acre. = *I. acreana*; nobil. = *I. nobilis*; vism. = *I. vismiifolia*.

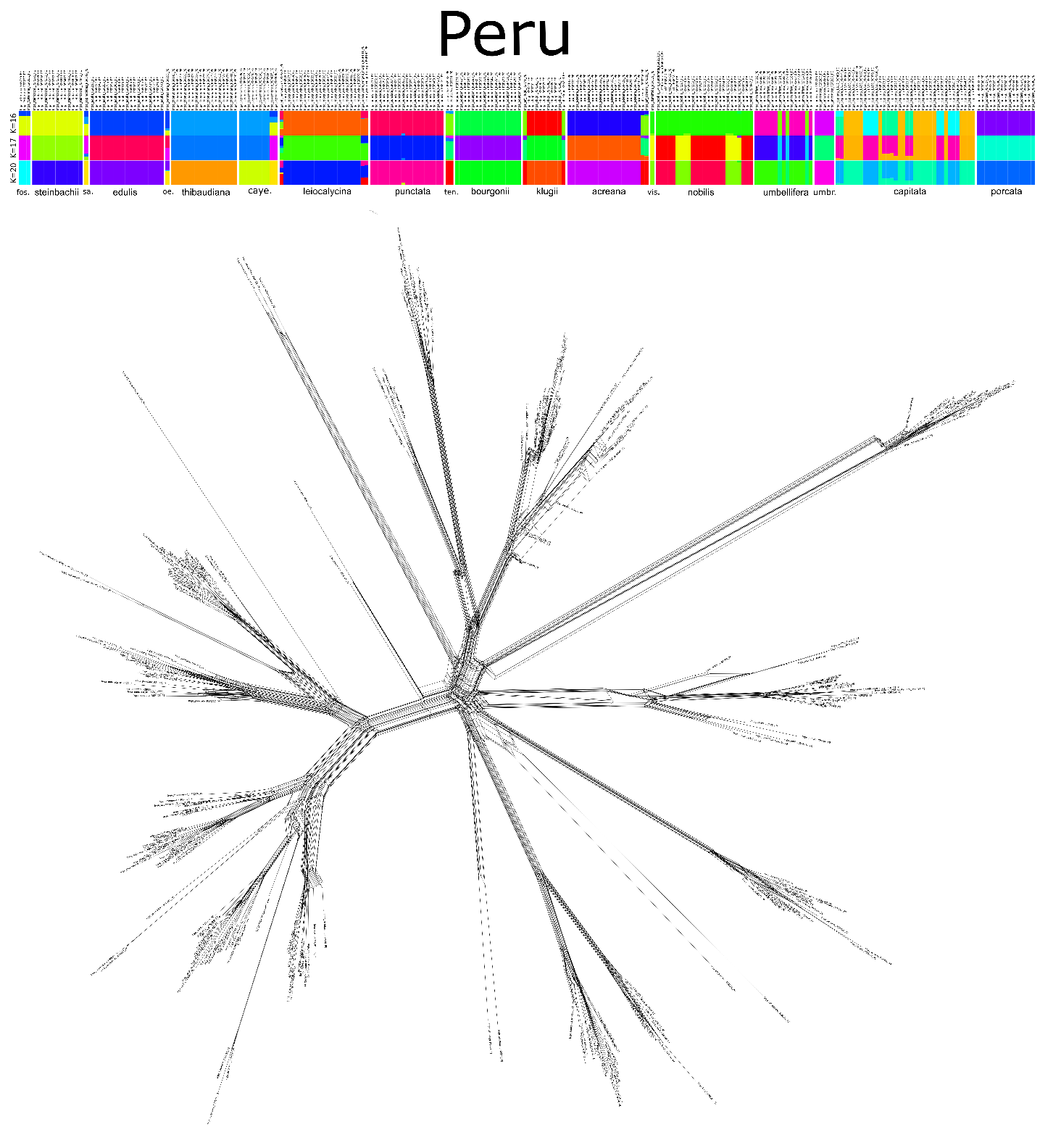

### Figure S2e. ADMIXTURE and SPLITSTREE population structure plots for Peruvian samples (see Figure S12 for high resolution)

### Figure 3a-3f. Interactive PCAs inferred for each regional community (a-e) as well as for all accessions together (f) based on whole-genome resequencing of all Inga accessions.

The zipped file contains six subfolders, corresponding to a PCA analysis for each regional community: Brazil (Figure S3a), CostaRica/Panama (Figure S3b), French Guiana (Figure S3c), Ecuador (Figure S3d), Peru (Figure S3e) and all samples together (Figure S3f). Each subfolder contains an interactive PCA and its accessory files (html), a PCA image (PDF) and a plot of variance explained by each principal component (PDF). In the PDF plots, the percent of variation explained by each PC is indicated under the x (PC1) and y (PC2) axes. The regional community from which an accession was collected is indicated by the shape of the point, whereas the morphological species identification given to the accession is marked by the colour of the point. Accessions that are more genetically similar cluster more closely together on PC1 and PC2.

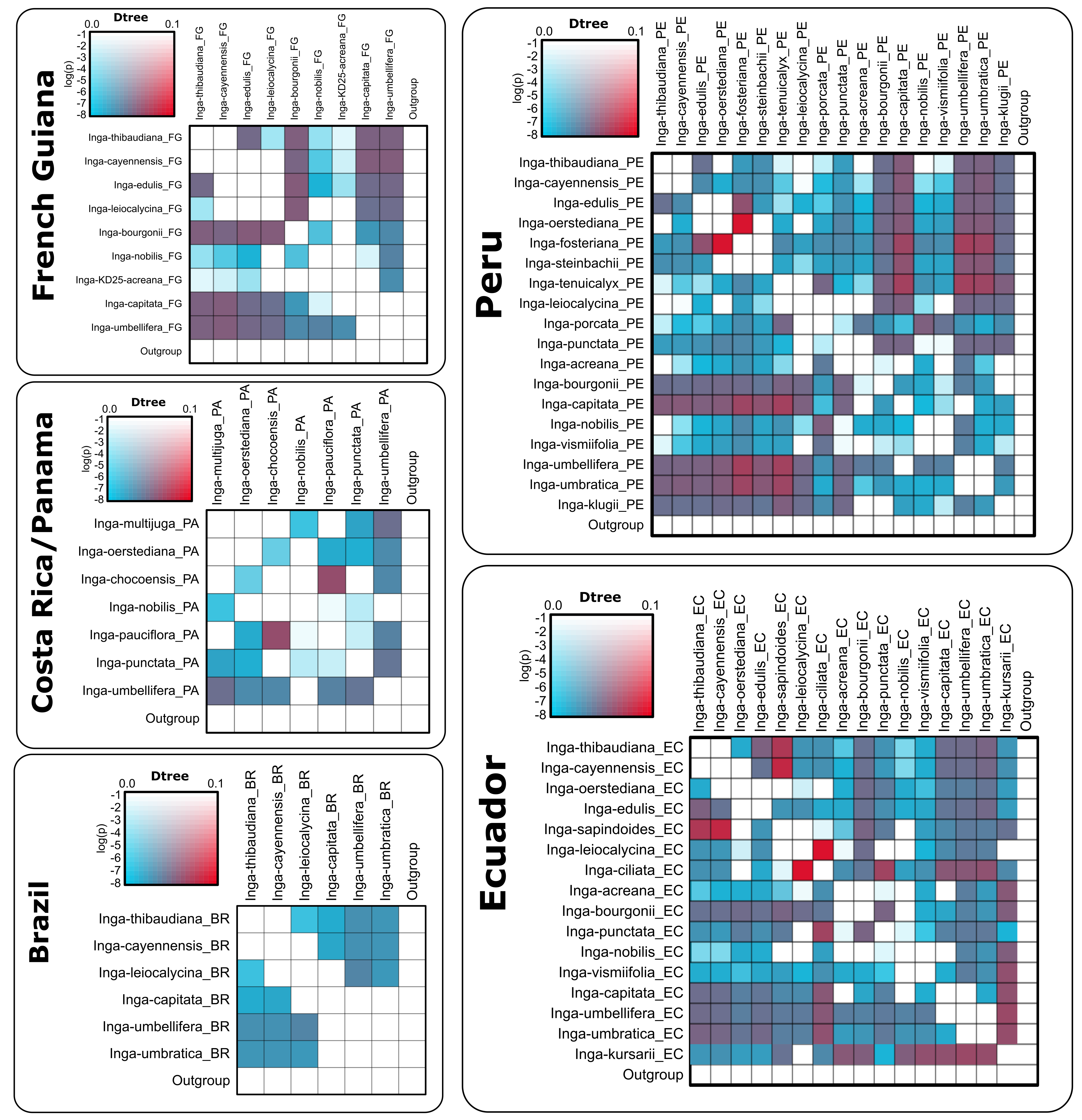

Figure S4. **Heatmaps of introgression intensity (D-statistic) for each regional community.**
Heatmaps of per-triplet D-statistic (D) indicating the degree of introgression between any two species, with 0 meaning no introgression and 1 indicating high introgression. Each heatmap was plotted for one of five regional *Inga* communities – Brazil, Costa Rica/Panama, French Guiana, Ecuador and Peru - based on whole-genome resequencing of accessions from each region. The colour of each square signifies the D-statistic (blue = low estimate; red = high estimate). The saturation of these colours represents the significance for that test (log(P)). D estimates were made from species trios ordered based on a species-level ‘guide tree’ inferred in phylogenetic analysis of the nuclear genome. This was to ensure that the P1/P2 taxa are more closely related to each other than to the P3 taxon and outgroup, as assumed by D-statistics.

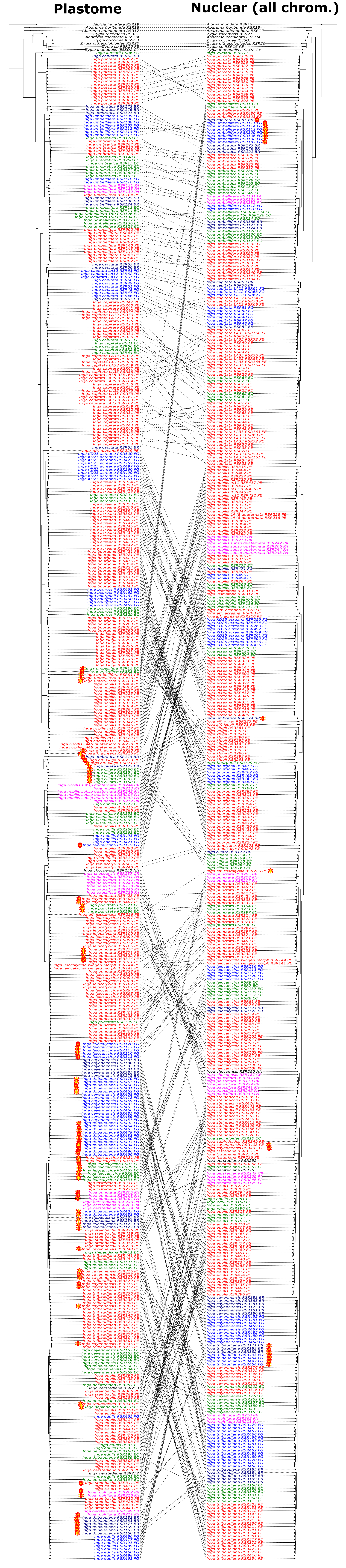

### Figure S5. Cophylo plot showing cytonuclear discordance between the plastome and nuclear genome (see Figure S13 for high resolution)

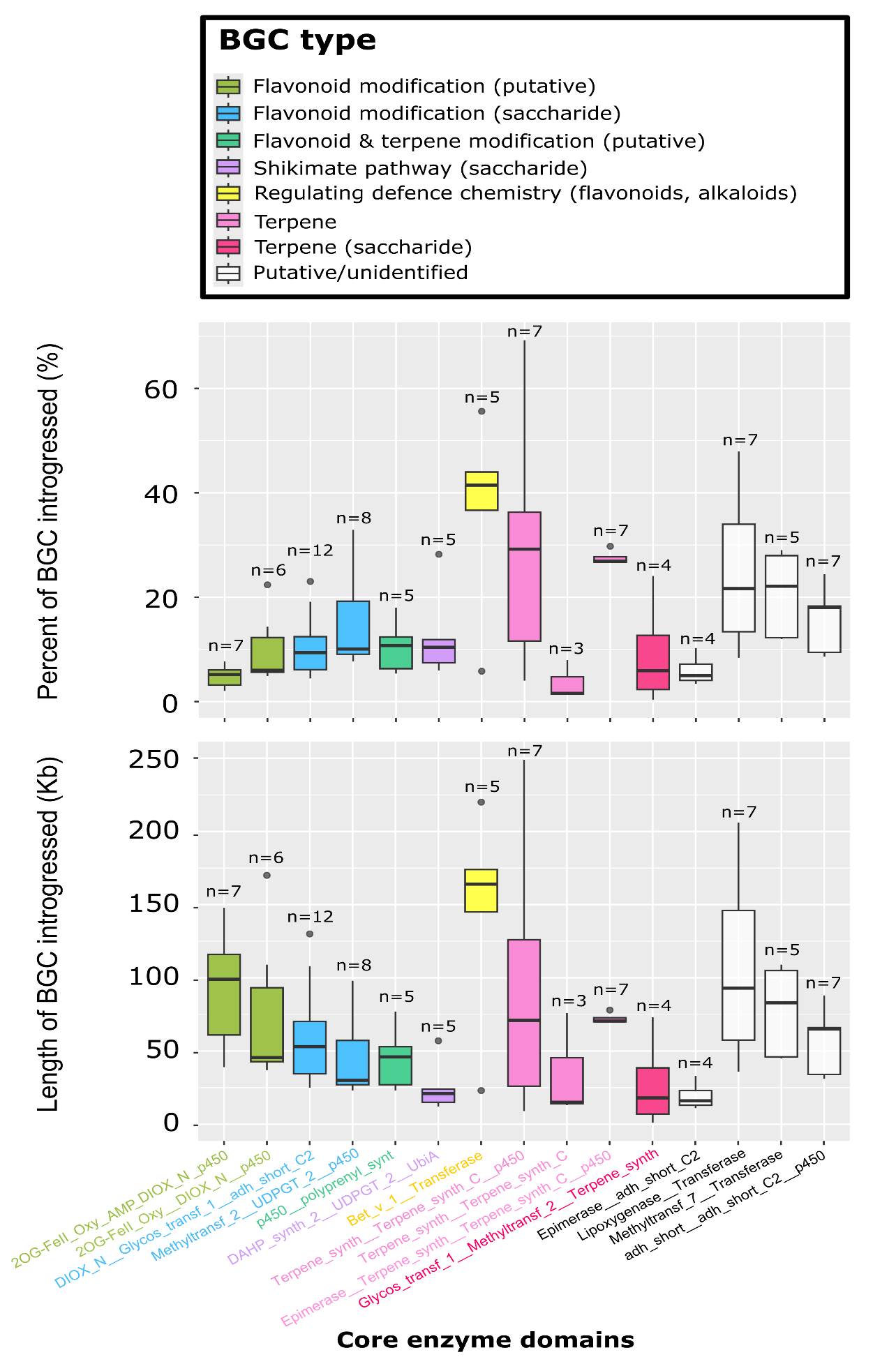

Figure S6. **Boxplots of introgressed block size for different Biosynthetic Gene Clusters.**
Box plots indicating the percentage (top panel) and length in Kilobases (bottom panel) of the longest contiguous blocks of introgressed variation we found for fifteen candidate biosynthetic gene clusters (BGCs) that had exceptionally high estimates of introgression, across a range of 23 *Inga* species pairs. Boxes are coloured by core enzyme domains that are coded for by the genes contained within each of the fifteen BGCs, which relates to the broad biosynthetic function with which they are associated (identified with PLANTISMASH). The number of species pairs in which we found introgression for each candidate BGC is labelled above each box. In the box plots, the dark bar represents the median, the top and bottom edges of each box represent the first and third quartiles, while the dark circles represent outliers.

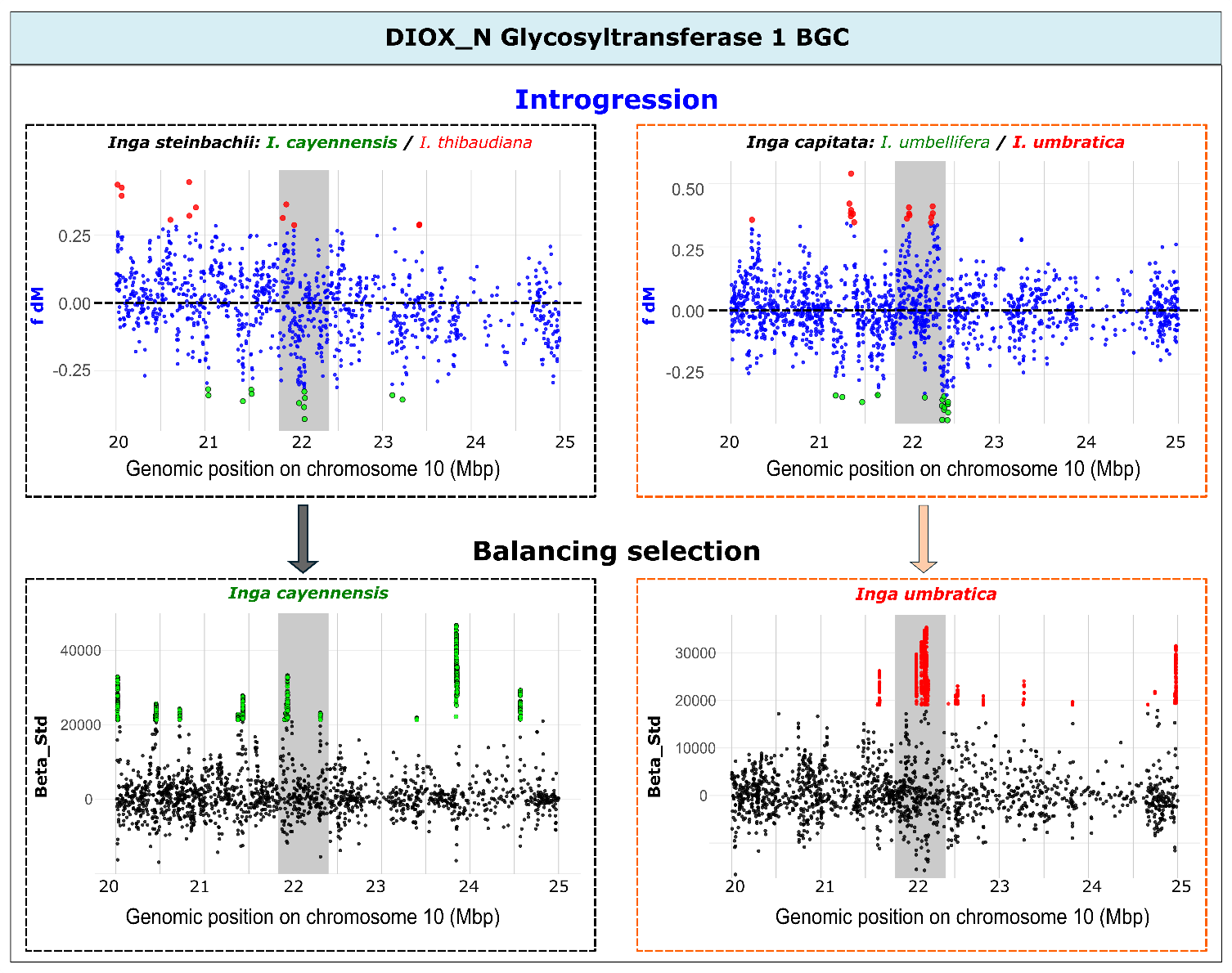

Figure S7. **Recurrent introgression and balancing selection in the same BGC between different species.**
Introgression scores (*f _dM_*: top two panels) and balancing selection scores (Beta_std: bottom two panels) in genomic windows surrounding the “*DIOX_N Glycosyltransferase_1”* biosynthetic gene cluster (BGC) on chromosome ten. The extent of the BGC is marked with a grey shaded box. For introgression scores, taxa used for each test are indicated above each plot, with P3 taxa in black, P1 taxa in green and P2 taxa in red. Positive *f _dM_* values (red outliers) indicate introgression between taxon P3 and P2, while negative scores (green outliers) indicate introgression between taxon P3 and P1. Outliers are first and 99th percentile values. For balancing selection scores, we plotted Beta_STD along the same region of the genome as for *f _dM_*, and plotted balancing selection scores for the taxon with the most overlap between Beta_STD and *f _dM_* outliers. Both 99th percentile outliers percentile and taxon names are coloured according to the corresponding P1 or P2 taxa in *f _dM_* analysis, with the chosen P1 or P2 taxon emboldened in the top two panels. Only positive Beta_STD values carry meaning, whereas negative values do not, and occur by chance when Watterson's theta is larger theta beta (see https://github.com/ksiewert/BetaScan). Different coloured dashed boxes indicate different sets of species with overlapping *f _dM_* and Beta_STD scores, both of which come from the Peru and Ecuador regional communities.

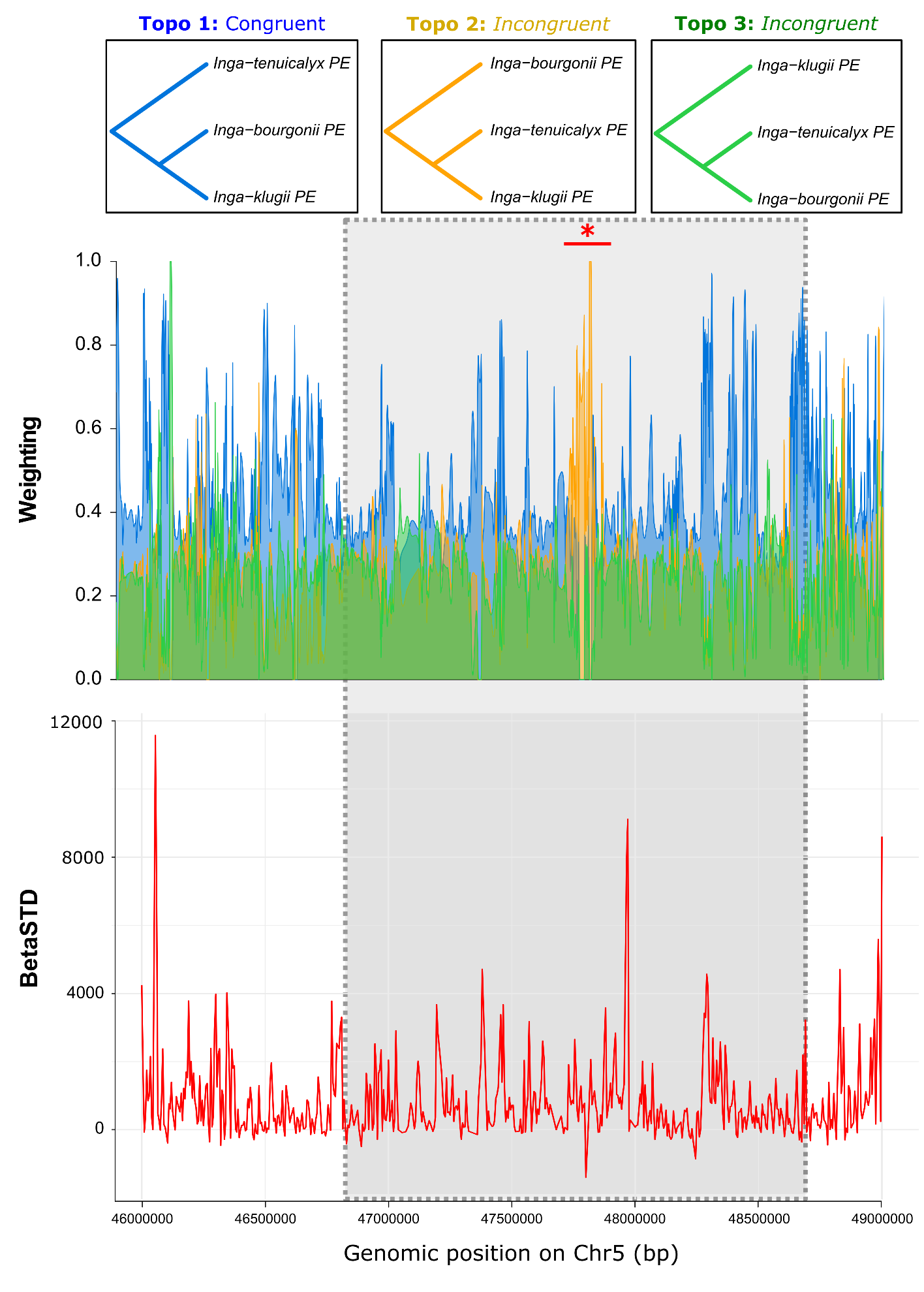

Figure S8. **TWISST2 plot of the longest introgressed gene block also experiencing balancing selection.**
TWISST2 plot, showing the longest introgressed gene block (top panel) that also experienced balancing selection (bottom panel). Specifically, the top panel shows a TWISST2 plot for the largest introgressed block of sequence within a candidate biosynthetic gene cluster (BGC), here indicating introgression between *Inga tenuicalyx* and *I. klugii* in Peru within the ‘2OG-FeII_Oxy, DIOX_N, p450 BGC’ plantiSMASH BGC (between 46841159-48756158 bp on chromosome five). The extent of the BGC is marked with a grey shaded box. The y-axis indicates topology weighting in 50kb smoothed windows along chromosome five. Topologies plotted above the main figure represent the three possible genealogies for the set of three taxa included in the TWISST2 analysis. Weighting of the most frequent topology, congruent with the species tree, is coloured in blue, whereas the two other incongruent topologies (which can arise from introgression, in which case two introgressing species are sister to one another) are coloured in yellow and green, respectively. Topology weighting corresponds to the frequency of each topology in a window.

The bottom panel shows standardised balancing selection scores (Beta_STD) in genomic windows surrounding the *“2OG-FeII_Oxy, DIOX_N, p450 BGC”* plantiSMASH BGC in *Inga klugii*. The extent of the BGC is marked with a grey shaded box. Only positive Beta_STD values carry meaning, whereas negative values do not, and occur by chance when Watterson's theta is larger theta beta (see https://github.com/ksiewert/BetaScan).

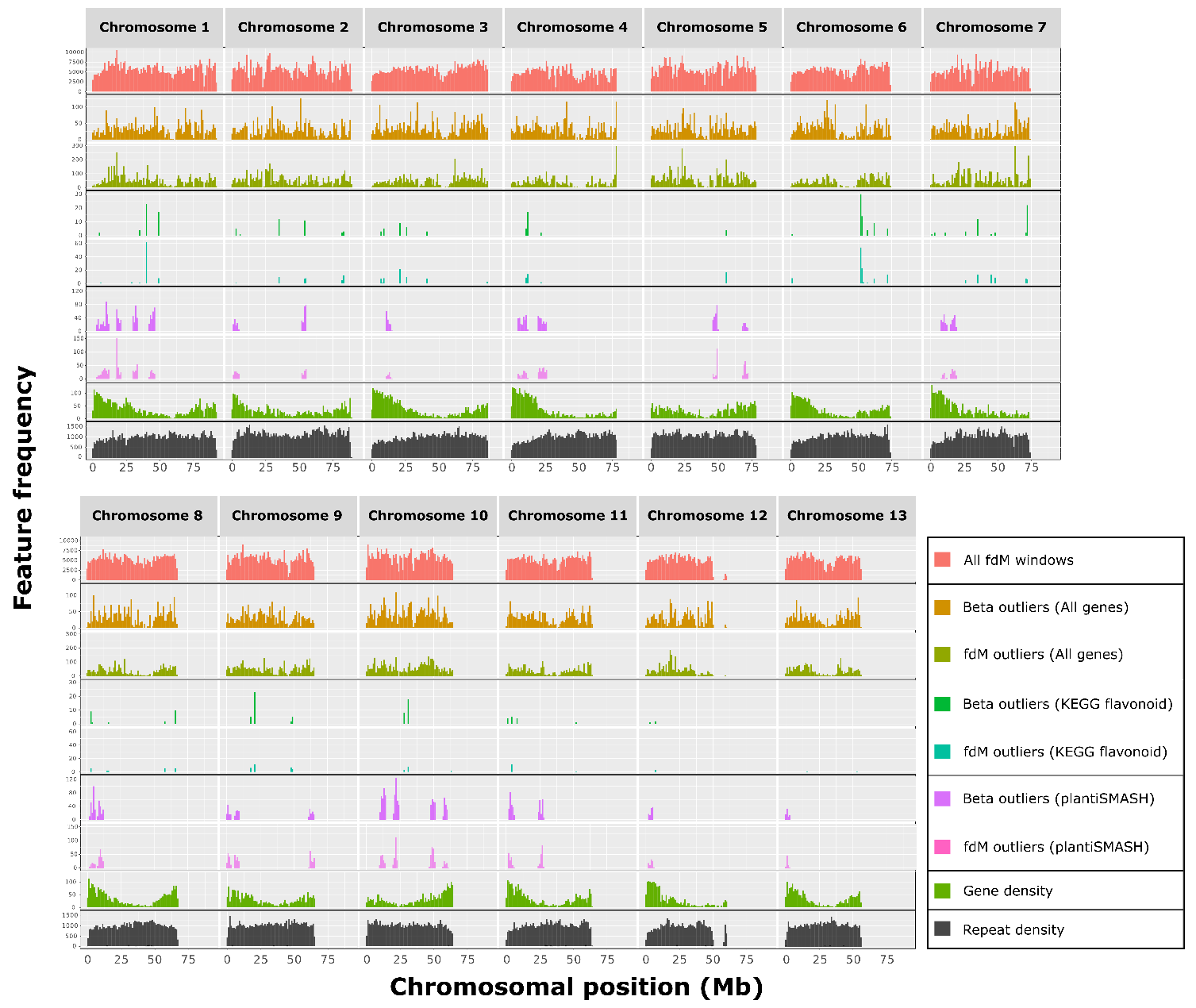

### Figure S9. Total count of different genomic features in focal accessions for introgression and balancing selection (see Figure S14 for high resolution).

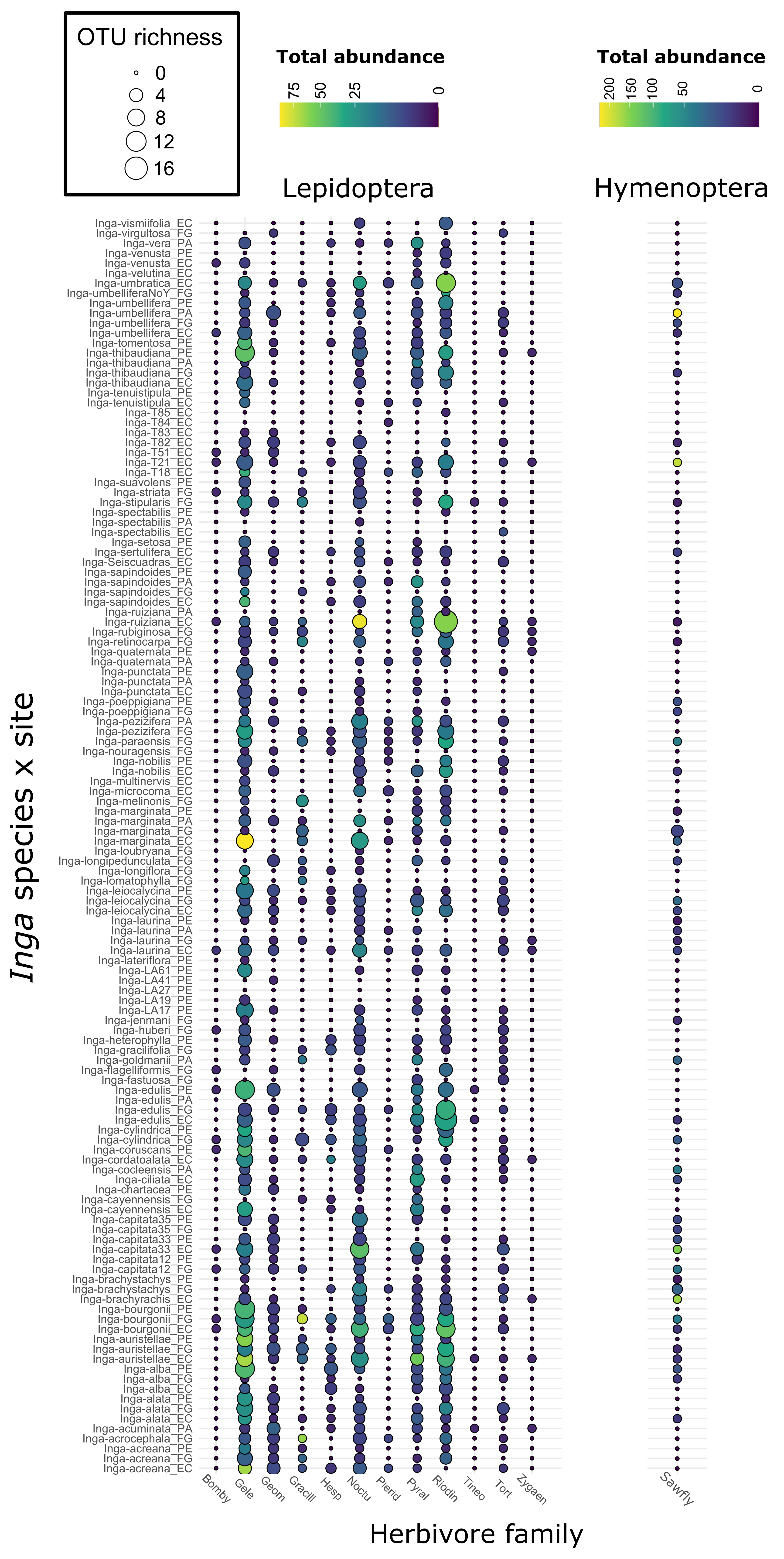

Figure S10. **Heatmap, indicating the structure of herbivore communities collected on different Inga species at different sites.**
Rows correspond to *Inga* species collected at each sampling site, and columns correspond to herbivore families. Circle colour represents the total abundance of herbivores within each herbivore family on each *Inga* species at a site (square-root transformed). Herbivore families are plotted according to insect order, each plotted on a different colour scale due to differing total herbivore abundances from each order (indicated in the legend above each order label). For both orders, circle size indicates OTU richness – the count of distinct herbivore OTUs within a herbivore family on an *Inga* species at a site. In herbivore family labels, shortened names correspond to: Bombycoidea (Bomby); Gelechiodea (Gele); Geometridae (Geom); Gracillariodea (Gracil); Hesperiidae (Hesp); Noctuoidea (Noctu); Pieridae (Pierid); Pyraloidea (Pyral); Riodinidae (Riodin); Tineoidea (Tineo); Torticidae (Tort); Zygaenoidea (Zygaen) and Argiidae (Sawfly). In *Inga* species x site labels, PE= Los Amigos, Perú; EC= Tiputiní, Ecuador; PA= Barro Colorado Island, Panama; FG= Nouragues, French Guiana

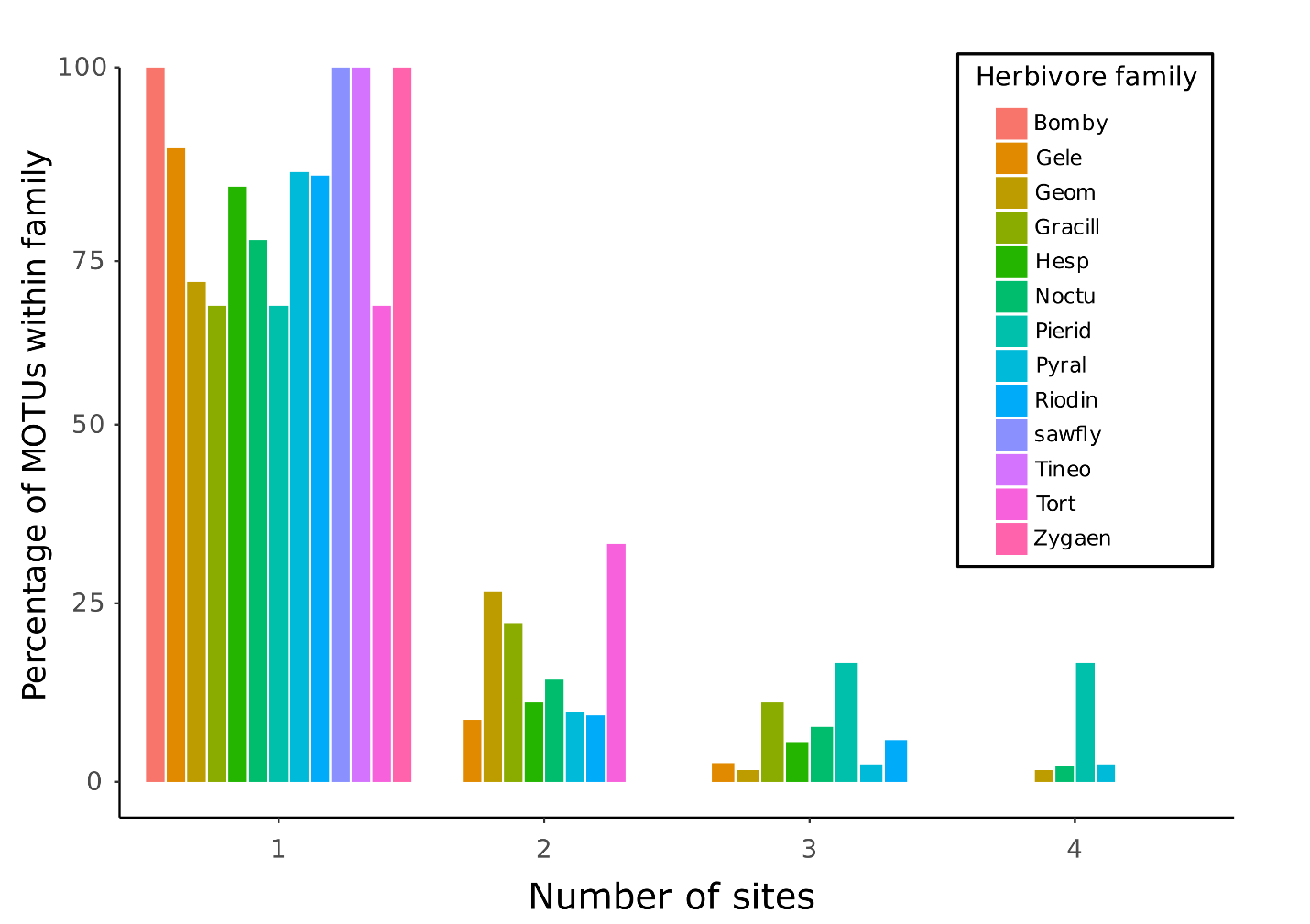

Figure S11. **Bar plot showing the percentage of herbivore MOTUs from each herbivore family found at differing numbers of sites.**
Bar plot showing the percentage of herbivore MOTUs from each herbivore family found at one site, two sites, three sites or all four sites (Los Amigos (Peru); Tiputiní (Ecuador); Barro Colorado Island (Panama) and Nouragues (French Guiana)). Bar colour indicates herbivore family – shortened herbivore family labels correspond to: Bombycoidea (Bomby); Gelechiodea (Gele); Geometridae (Geom); Gracillariodea (Gracil); Hesperiidae (Hesp); Noctuoidea (Noctu); Pieridae (Pierid); Pyraloidea (Pyral); Riodinidae (Riodin); Tineoidea (Tineo); Torticidae (Tort); Zygaenoidea (Zygaen) and Argiidae (Sawfly).

Captions of Extended Data Figures in separate files

### Fig 3a-3f: Interactive PCAs inferred for each regional community (a-e) as well as for all accessions together (f) based on whole-genome resequencing of all Inga accessions.

Interactive PCAs inferred for each regional community (a-e) as well as for all accessions together (f) based on whole-genome resequencing of all *Inga* accessions. The zipped file contains six subfolders, corresponding to a PCA analysis for each regional community: Brazil (Figure S3a), CostaRica/Panama (Figure S3b), French Guiana (Figure S3c), Ecuador (Figure S3d), Peru (Figure S3e) and all samples together (Figure S3f). Each subfolder contains an interactive PCA and its accessory files (html), a PCA image (PDF) and a plot of variance explained by each principal component (PDF). In the PDF plots, the percent of variation explained by each PC is indicated under the x (PC1) and y (PC2) axes. The regional community from which an accession was collected is indicated by the shape of the point, whereas the morphological species identification given to the accession is marked by the colour of the point. Accessions that are more genetically similar cluster more closely together on PC1 and PC2.

### Figure S12. ADMIXTURE and SPLITSTREE population structure plots for Peruvian samples

The top panel shows *ADMIXTURE* plot generated using the K values (i.e., the number of estimated genetic clusters) with the lowest cross-validation errors. The *ADMIXTURE* plot indicates ancestry proportions from *K* inferred genetic groups, which are indicated with different colours. Each column represents one accession, with accession names above the column. Morphological species identifications for groups of accessions are labelled below the x axis. The bottom panel shows a *SplitsTree* built using uncorrelated P distances, where connecting edges indicate shared variation. Both plots were inferred using whole-genome resequencing data for *Inga* accessions collected from Peru. In the ADMIXTURE plot labels, fos.=*I. fosteriana*; sa.= *I. sapindoides*; oe. = *I. oerstediana;* caye. = *I. cayennensis; ten.* = *I.tenuicalyx; vis.* = *I. vismiifolia ; umbr.* = *I. umbratica.*

### Figure S13. Cophylo plot showing cytonuclear discordance between the plastome and nuclear genome.

Phylogenetic trees inferred with IQtree for the nuclear genome (randomly downsampled SNPs from across all chromosomes) and the plastome (all SNPs) for all sequenced accessions. Disagreements between tree topologies are indicated with crossed lines connecting the same accession between the two trees. Cases where accessions do not group with their conspecifics are marked by a yellow star in both trees.

### Figure S14. Total count of different genomic features in focal accessions for introgression and balancing selection

Total count of different genomic features across all accessions used in the balancing selection (betaSTD) and introgression (*f _dM_*) analyses, counted in 1 megabase (Mb) windows across the genome. From top to bottom, the first column indicates the total number of *f _dM_* windows in a 1Mb length of the genome, given that *f _dM_* windows are based on having 50 informative SNPs and so may vary in absolute numbers of base pairs. The second to ninth columns indicate the total count of 99^th^ percentile outliers for balancing selection (Beta) and introgression (*f _dM_*) found within each class of genomic annotation: all coding genes, all KEGG flavonoid genes and all plantiSMASH BGCs. These outliers were counted across all runs of *f _dM_* and BetaScan. The final two columns indicate the total density of coding genes and the total density repetitive elements in each 1MB window of the *Inga leiocalycina* reference genome, to which all whole-genome resequencing data were mapped. Chromosome number is indicated above each column, with the position on each chromosome indicated below each column.

Extended Data Tables (available as separate files)

Table S1.
Accession information for whole genome resequencing samples included in this study. Shown are sample accession identifiers, collector names and numbers, country, region, and locality of collection. Notes indicate habitat type (e.g., floodplain, terra firme), sampling location information or associated chemotype information where available.

Table S2.
Sample inclusion and population assignments for analyses of introgression (Dsuite, including D statistics, F_4_ ratios, F_branch_, *f _dM_*) and balancing selection (BetaScan2). For each accession, we report population assignment and regional community of origin, and indicate whether the sample was included in the Dsuite introgression analysis and/or the BetaScan balancing selection analysis. “YES” denotes inclusion in the corresponding analysis.

Table S3.
Explanations for population trios and pairwise comparisons selected for per-locus introgression analyses (*f _dM_*) and demographic inference of long-term barriers to gene flow (gIMble).

**a)** *f _dM_*: For each trio, we list the focal species (P1–P3), the biological rationale for inclusion (e.g., introgression signal inferred from F_branch_ statistics), qualitative similarity of major defensive chemistry among taxa based on *50*.

**b)** gIMble: Species pairs used for demographic modelling, indicating whether pairs differ in defensive chemistry, whether they are sister species, and whether they were additionally included in the *f _dM_* analysis.

Table S4.
Cross-validation results for ancestry model selection in ADMIXTURE. Shown are cross-validation (CV) errors for ancestry inference models run with the data and varying numbers of genetic clusters (K) for each of the regional communities (Brazil, CostaRica/Panama, French Guiana, Ecuador and Peru). Lower CV error indicates better model support. CV values are plotted against number of genetic clusters (K) to assess the best value of K in each analysis. The best K value, indicated by the lowers CV error value, is emboldened and highlighted in green. Where lower CV errors are very close together (within 0.01 of each other), the next-lowest value is emboldened and highlighted in yellow. Highlighted K values are plotted in the final ADMIXTURE plots (**Figure** S2a-S2e).

Table S5.
Number of barrier windows with reduced gene flow inferred by gIMble for each species pair studied. Columns from left to right denote: (1-3) Species pair (labelled species A and B), region from which they were sampled and demographic model used for inferences of blockwise gene flow, which were selected following comparisons between multiple different models; (4) Total count of genomic windows used in the analysis; (5) Total number of windows with deltaB <0, denoting that a demographic model without gene flow fits best, suggesting a window is a barrier to gene flow; (6) The percentage of all windows that were inferred to be barriers to gene flow.

Table S6.
Biosynthetic gene clusters (BGCs) identified by PlantiSMASH using the *Inga leiocalycina* reference genome. For each cluster, we report the predicted gene cluster type, cluster identifier, chromosomal location and coordinates, physical size, core biosynthetic protein domains, number of CD-HIT clusters, total number of genes, and the gene and protein accessions comprising each cluster. Putative functional annotations are based on conserved domain content.

Table S7.
Fine-scale overlap in per-window *f _dM_* introgression outliers and annotated genomic features.

**a)** *f _dM_* outlier regions overlapping plantiSMASH-predicted biosynthetic gene clusters. For each comparison, we report the species pair, overlapping plantiSMASH cluster, chromosomal location and size, gene cluster type and core biosynthetic domains, midpoint of the PlantiSMASH BGC location (*ps_mid*), the distance between the midpoints of the *f _dM_* introgression window and the PlantiSMASH BGC (*hit_dist*), *f _dM_* introgression value and midpoint (*f _dM_*_mid), and whether the *f _dM_* window midpoint falls within a plantiSMASH-defined cluster.

**b)** Individual genes overlapping *f _dM_* introgression window. Shown are the species pair, gene coordinates and annotations, gene midpoint (*gff*_mid), the distance between the midpoints of the *f _dM_* introgression window and the PlantiSMASH BGC (*hit_dist*), *f _dM_* introgression value and midpoint (*f _dM_*_mid), and whether the gene overlaps an introgressed region.

Table S8.
TWISST2 results, showing introgressed block sizes for each candidate BGC in each species trio.

**a)** Summary of the longest introgressed block, inferred by TWISST2, for each candidate BGC per candidate species pair – species pairs were selected based on outlying introgression estimates in the BGC. For each run, we report whether a candidate BGC/species pair combination contained both introgression (*f _dM_*) and balancing selection (BetaSTD) outliers, or just introgression outliers. We also report the plantiSMASH BGC ID, gene cluster type, core biosynthetic domains, chromosome, genomic coordinates, total size, and number of individual genes found in each candidate plantiSMASH BGC. The next column shows the conflicting genealogical topology output from TWISST2 that indicated introgression between the candidate species pair. Next are the largest introgressed block size in megabases (Mb) within the candidate BGC, the percentage of the BGC encompassed by introgressed sequence, the genomic coordinates of the largest introgressed block, and the number of genes in the largest introgressed block. For each candidate BGC/species pair combination, we also list the mean and standard deviation of the longest introgressed block in a random draw of 10 regions of similar size to each BGC on the same chromosome, and whether the largest block of introgressed sequence in the BGC was larger than that random draw. The final three columns denote the species pair implicated in introgression and the highest fdM score within the same candidate BGC for those two species.

**b)** Longest introgressed block length for each topology in each candidate BGC/species pair analysis in TWISST2. In the ‘Topology’ column, the most frequent genealogical topology is congruent with the species tree and is denoted “Topo1”. The two other incongruent topologies (which can arise from introgression, in which case two introgressing species are sister to one another) are denoted as “Topo2” and “Topo3”. Genomic coordinates and total length of the longest block found for each topology within each candidate BGC/species pair combination are listed in the subsequent three columns.

**c)** Longest introgressed block length for each topology among ten random draws, each of which were of the same length and on the same chromosome as the corresponding candidate BGC in each candidate species pair. In the ‘Topology’ column, the most frequent genealogical topology is congruent with the species tree and is denoted “Topo1”. The two other incongruent topologies (which can arise from introgression, in which case two introgressing species are sister to one another) are denoted as “Topo2” and “Topo3”. Genomic coordinates and total length of the longest block found for each topology within each candidate BGC/species pair combination are listed in the subsequent three columns.

Table S9.
Fine-scale overlap in per-window BetaSTD balancing selection outliers and annotated genomic features.

**a)** *BetaSTD* outlier regions overlapping plantiSMASH-predicted biosynthetic gene clusters. For each comparison, we report the population analysed (in the format Inga-species_Country, where PE = Peru, EC = Ecuador, FG = French Guiana, BR = Brazil and PA = Panama). We also report overlapping plantiSMASH cluster, chromosomal location and size, gene cluster type and core biosynthetic domains, midpoint of the PlantiSMASH BGC location (*ps_mid*), the distance between the midpoints of the *BetaSTD* balancing selection window and the PlantiSMASH BGC (*hit_dist*), *BetaSTD* balancing selection value and midpoint aggregated into standardised 5kb windows (*Beta1_std_agg*), and whether the *BetaSTD* window midpoint falls within a plantiSMASH-defined cluster.

**b)** Individual genes overlapping *BetaSTD* balancing selection window. Shown are the analysed population, gene coordinates and annotations, gene midpoint (*gff*_mid), the distance between the midpoints of the *BetaSTD* balancing selection window and the PlantiSMASH BGC (*hit_dist*), *BetaSTD* balancing selection value and midpoint aggregated into standardised 5kb windows (*Beta1_std_agg*), and whether the *BetaSTD* window midpoint falls within a plantiSMASH-defined cluster.

Table S10.
Fine-scale overlap between per-window BetaSTD balancing selection outliers and *f _dM_* introgression outliers within the same species, and annotated genomic features.

**a)** Overlapping *BetaSTD* and *f _dM_* outlier regions with plantiSMASH-predicted biosynthetic gene clusters. We report the overlapping plantiSMASH cluster name, chromosomal location and size, gene cluster type and core biosynthetic domains and midpoint of the PlantiSMASH BGC location (*ps_mid*). For balancing selection outliers, we report the population analysed (in the format Inga-species_Country, where PE = Peru, EC = Ecuador, FG = French Guiana, BR = Brazil and PA = Panama). We also report the distance between the midpoints of the *BetaSTD* balancing selection window and the PlantiSMASH BGC (*hit_dist_betaSTD*), *BetaSTD* balancing selection value and midpoint aggregated into standardised 5kb windows (*Beta1_std_agg*). For introgression outliers overlapping the same plantiSMASH BGC as the balancing selection outlier in the same row, we report the species pair (one of which will be identical to the population analysed for BetaSTD), the distance between the midpoints of the *f _dM_* introgression window and the PlantiSMASH BGC (*hit_dist_fdM*), *f _dM_* introgression value and midpoint (*f _dM_*_mid).

**b)** Overlapping *BetaSTD* and *f _dM_* outlier regions with individual genes. We report the overlapping gene ID, chromosomal location and midpoint of the gene (*gff_mid*). For balancing selection outliers, we report the population analysed, alongside the distance between the midpoints of the *BetaSTD* balancing selection window and the gene (*hit_dist_betaSTD*), *BetaSTD* balancing selection value and midpoint aggregated into standardised 5kb windows (*Beta1_std_agg*). For introgression outliers overlapping the same gene as the balancing selection outlier in the same row, we report the species pair (one of which will be identical to the population analysed for BetaSTD), the distance between the midpoints of the *f _dM_* introgression window and the gene (*hit_dist_fdM*), *f _dM_* introgression value and midpoint (*f _dM_*_mid).

Table S11.
Partitioning of variation in regional herbivore community composition among geographic regions and host *Inga* species, estimated using Bray-Curtis Distances. Results from a permutational analysis of variance (PermANOVA) testing the effects of regional herbivore community across all herbivore species, with three models: one assessing how *Inga* host species influences herbivore turnover, one assessing how regional community affects herbivore turnover, and one assessing how host *Inga* + regional community affects herbivore turnover. Total sample size (N) is shown under each model name. Columns in the table refer to the model type, the degrees of freedom for each parameter in each model (Summary and DF), sums of squares, variance explained (R²), F statistics and permutation-based *P*-values. Significance was assessed using permutation tests with 99999 permutations. Significant *P*-values (P<0.05) are emboldened.

Table S12.
Effects of geographic site and host *Inga* species on herbivore community dissimilarity (phylogenetic turnover, estimated with weighted UniFrac, dw) within major herbivore families. PermANOVA results are grouped for each herbivore family, each testing three models: one assessing how *Inga* host species influences herbivore phylogenetic turnover, one assessing how regional community affects herbivore phylogenetic turnover, and one assessing how host *Inga* + regional community affects herbivore phylogenetic turnover. Total sample size (N) is shown under each model name. Columns in the table refer to the herbivore family assessed, model type, the degrees of freedom for each parameter in each model (Summary and DF), sums of squares, variance explained (R²), F statistics, alongside permutation-based *P*-values and Bonferroni-corrected *P*-values. Significance was assessed using permutation tests with 99999 permutations. Significant P-values (P<0.05) are emboldened.

Table S13.
Mantel tests relating herbivore community similarity (estimated with Bray-Curtis distances) to similarity in balancing selection profiles within PlantiSMASH BGCs and KEGG Flavonoid genes among different *Inga* populations. Mantel statistic (R), permutation-based *P-*values, number of permutations and the total number of *Inga* populations (N) involved in each model are shown. Significant associations (P<0.05) are emboldened.

Table S14.
Herbivore family-specific Mantel tests, testing the association between herbivore community dissimilarity (phylogenetic turnover, estimated with weighted UniFrac, dw) and similarity in balancing selection profiles within PlantiSMASH BGCs and KEGG Flavonoid genes among different *Inga* populations. Mantel tests were performed per herbivore-family (denoted in column “Herbivore_family”). Mantel statistic (R), permutation-based *P*-values, Bonferroni-adjusted *P-*values, number of permutations, and the total number of *Inga* populations (N) involved in each model are reported in the table, with significant associations (P<0.05) emboldened.
