## Supplementary figures and images for "Hybridisation and herbivory fuel Amazonian tree radiations"

### Figure S12

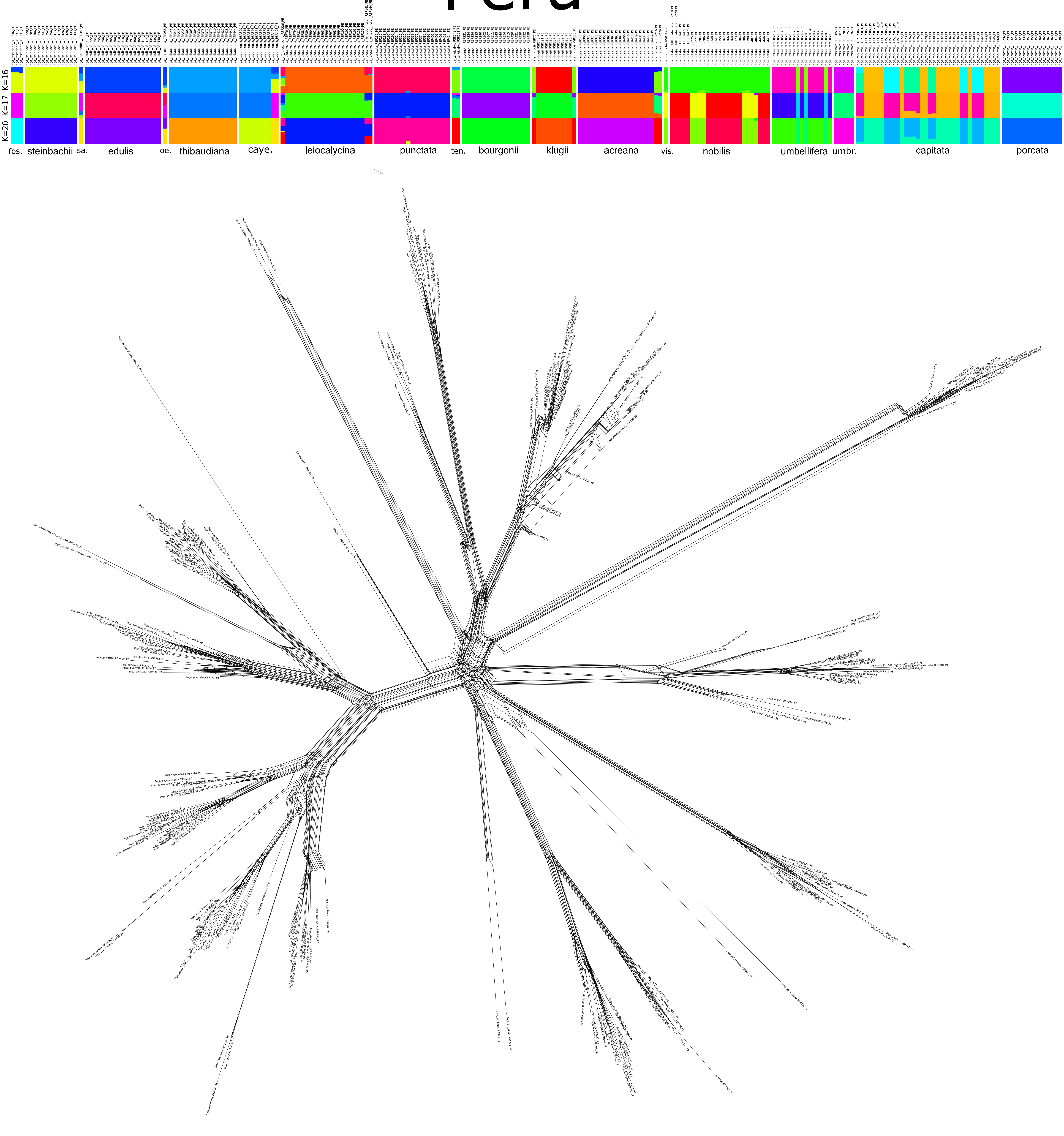

### Figure S13

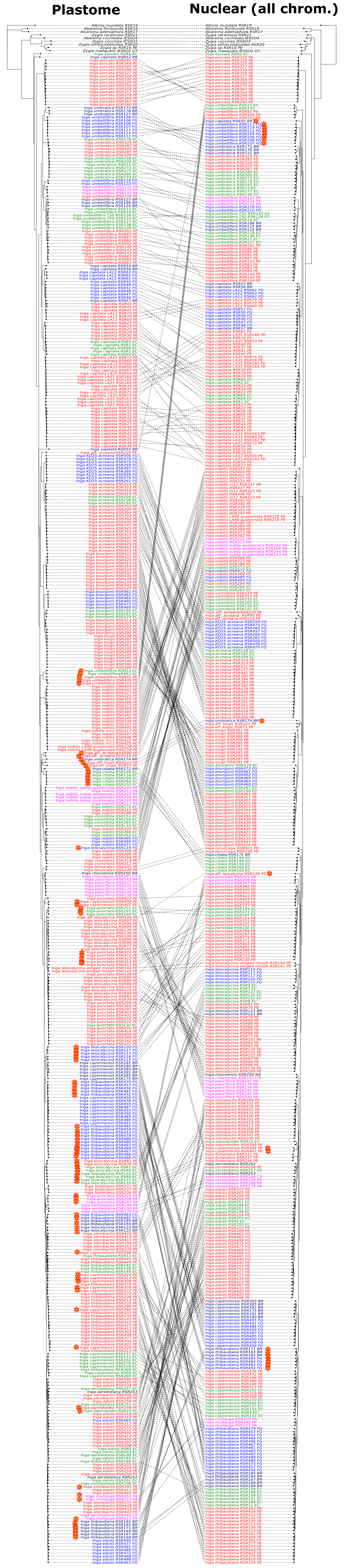

### Figure S14

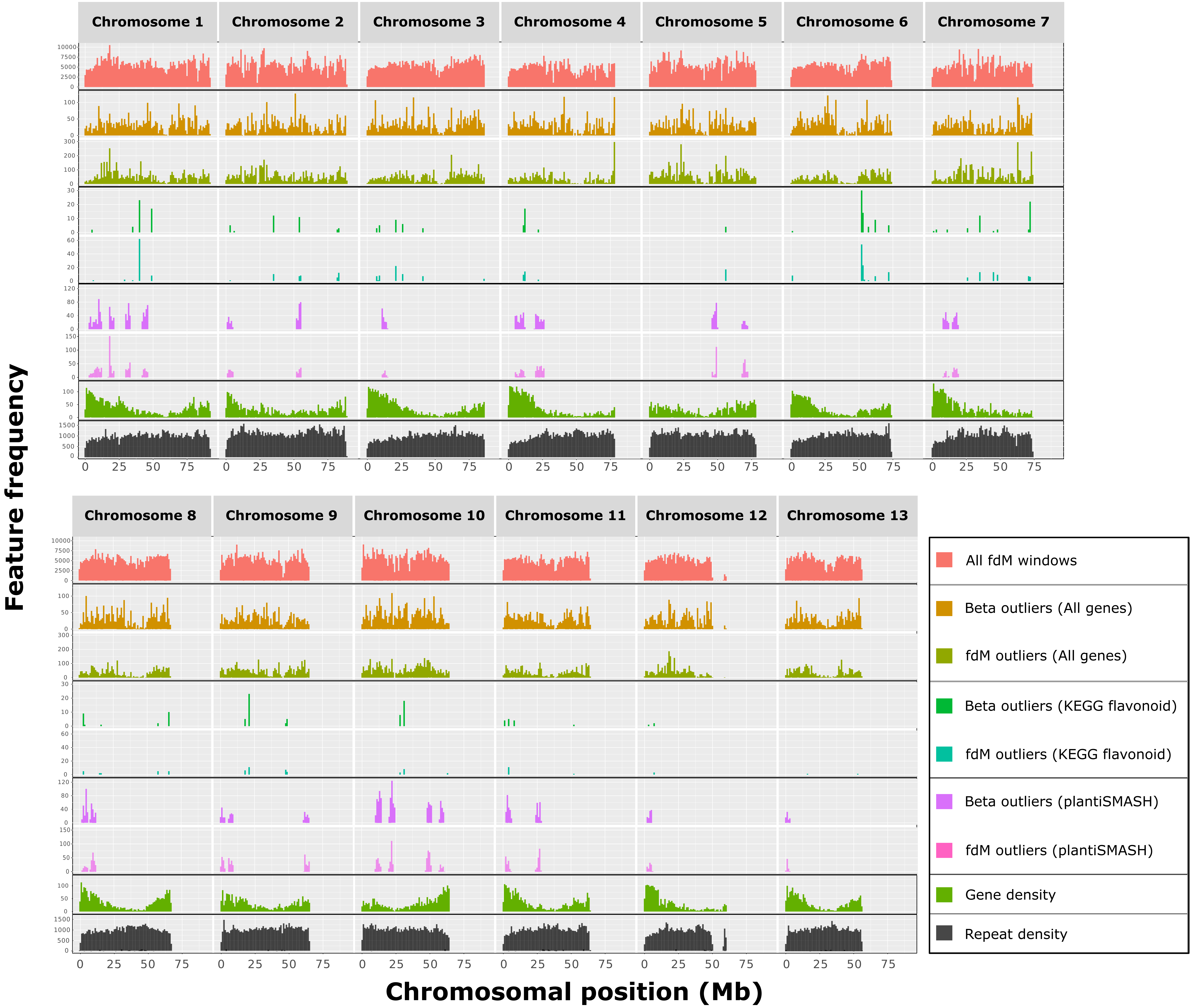
